# Seed Genome Hypomethylated Regions Are Enriched In Transcription Factor Genes

**DOI:** 10.1101/356501

**Authors:** Min Chen, Jer-Young Lin, Jungim Hur, Julie M. Pelletier, Russell Baden, Matteo Pellegrini, John J. Harada, Robert B. Goldberg

## Abstract

The precise mechanisms that control gene activity during seed development remain largely unknown. Previously, we showed that several genes essential for seed development, including those encoding storage proteins, fatty acid biosynthesis enzymes, and transcriptional regulators, such as *ABI3* and *FUS3*, are located within hypomethylated regions of the soybean genome. These hypomethylated regions are similar to the DNA methylation valleys (DMVs), or canyons, found in mammalian cells. Here, we address the question of the extent to which DMVs are present within seed genomes, and what role they might play in seed development. We scanned soybean and *Arabidopsis* seed genomes from post-fertilization through dormancy and germination for regions that contain < 5% or < 0.4% bulk methylation in CG-, CHG-, and CHH-contexts over all developmental stages. We found that DMVs represent extensive portions of seed genomes, range in size from 5 to 76 kb, are scattered throughout all chromosomes, and are hypomethylated throughout the plant life cycle. Significantly, DMVs are enriched greatly in transcription factor genes, and other developmental genes, that play critical roles in seed formation. Many DMV genes are regulated with respect to seed stage, region, and tissue - and contain H3K4me3, H3K27me3, or bivalent marks that fluctuate during development. Our results indicate that DMVs are a unique regulatory feature of both plant and animal genomes, and that a large number of seed genes are regulated in the absence of methylation changes during development - probably by the action of specific transcription factors and epigenetic events at the chromatin level.

**Significance:** We scanned soybean and *Arabidopsis* seed genomes for hypomethylated regions, or DNA Methylation Valleys (DMVs), present in mammalian cells. A significant fraction of seed genomes contain DMV regions that have < 5% bulk DNA methylation, or, in many cases, no detectable DNA methylation. Methylation levels of seed DMVs do not vary detectably during seed development with respect to time, region, and tissue, and are present prior to fertilization. Seed DMVs are enriched in transcription factor genes and other genes critical for seed development, and are also decorated with histone marks that fluctuate with developmental stage, resembling in significant ways their animal counterparts. We conclude that many genes playing important roles in seed formation are regulated in the absence of detectable DNA methylation events, and suggest that selective action of transcriptional activators and repressors, as well as chromatin epigenetic events play important roles in making a seed - particularly embryo formation.

## Introduction

Seeds are the agents for higher plant sexual reproduction, and an essential source of food for human and animal consumption. They consist of three major regions - the seed coat, endosperm, and embryo - which have different genetic origins, unique functions, and distinct developmental pathways (1). The seed coat transfers nutrients from the maternal plant to the embryo during seed development, and protects the seed during dormancy - a period of quiescence where growth and development have ceased temporarily. The endosperm also provides nourishment to the embryo, particularly during early embryogenesis, and in dicots, degenerates and remains as a vestigial cell layer in the mature seed (2). The embryo, on the other hand, differentiates into axis and cotyledon regions - the former giving rise to the mature plant after germination, while the latter is terminally differentiated and accumulates storage reserves that are used as an energy source for the germinating seedling (1, 3). Gene activity is highly regulated with respect to space and time within each seed region (4). However, the detailed mechanisms required for the differentiation of each seed region remain to be identified.

Selective DNA methylation of maternal and paternal alleles, or imprinting, plays a critical role in endosperm development (5-7). Imprinting, however, does not appear to play a major role in embryo formation (6, 7). The extent to which DNA methylation events regulate the activity of specific genes during embryogenesis is largely unknown. Recently, we (8), and others (9-11), showed that, on a global basis, CG- and CHG-context methylation does not change significantly during seed development. By contrast, CHH-context methylation increases primarily within transposons during the period leading up to dormancy (8-11), and appears to be a fail-safe mechanism to ensure that transposons remain silent and do not inactivate genes essential for seed development and germination (8).

In the course of our seed methylome studies with both soybean and *Arabidopsis*, we identified several genes important for seed formation, including storage protein and fatty acid metabolism genes, which are located within hypomethylated genomic regions that remain static with respect to DNA methylation throughout development (8). These regions resemble the DNA methylation valleys (DMVs) (12), hypomethylated canyons (UMRs) (13), and non-methylated islands (NMIs) (14) found in many animal cell types which are enriched with transcription factor genes and coated with specific histone marks. Genes within these DMVs are not regulated by DNA methylation events, but by both transcriptional and epigenetic processes at the chromatin level (12-15).

In this study, we scanned soybean and *Arabidopsis* seed genomes for DMVs. We found that a significant fraction of these seed genomes contain DMV regions that have < 5% bulk DNA methylation, or, in many cases, no detectable DNA methylation. Methylation levels of seed DMVs do not vary detectably during seed development with respect to time, region, and tissue, and appear to present prior to fertilization. Seed DMVs are enriched in transcription factor genes and other genes critical for seed development, and are also decorated with histone marks that fluctuate with developmental stage, resembling in significant ways their animal counterparts. We conclude that many genes playing important roles in seed formation are regulated in the absence of detectable DNA methylation events, and suggest that selective action of transcriptional activators and repressors, as well as chromatin epigenetic events play important roles in making a seed - particularly embryo formation.

## Results

**DMVs Represent a Significant Portion of Soybean Seed Genomes**. We scanned soybean seed methylomes from the globular stage through dormancy and germination for regions with < 5% bulk methylation in all cytosine contexts (CG, CHG, and CHH) using a sliding 5 kb window with 1 kb smaller steps to search for hypomethylated regions, or DMVs (Figs. 1 and 2*A*). Seed DMVs were defined as genomic regions with < 5% bulk methylation over all developmental stages investigated (see *Materials and Methods*). This strategy identified 21,669 seed DNA regions, or 21% of the soybean genome (210 Mb), that were hypomethylated and did not vary significantly throughout seed development, or during early seed germination, with respect to methylation status (Fig. 2B, Table 1, and Dataset S1). Ninety-nine percent of seed DMVs identified during seed development was also shared with seedling and seedling cotyledon DMV regions, and, as such, we refer to all DMVs uncovered by our genome scans as seed DMVs (see *Materials and Methods*). Detailed analysis showed that (i) the majority of seed DMVs had bulk methylation levels between 0 to 1% in all cytosine contexts (*Appendix Fig. S1A*), and (ii) the average bulk methylation level per DMV was 0.14% (*Appendix Fig. S1B*), or, on average, one methylated cytosine per 1 kb of DMV (*Appendix Fig. S1B*), indicating that our strategy was robust enough to identify genomic regions that were significantly hypomethylated over all developmental stages. By contrast, the average bulk methylation level of the soybean genome was 11.5%, or, on average, 43 methylated cytosines per 1 kb (*Appendix Fig. S1B*). We validated this approach by scanning soybean seed methylomes at a < 0.4% bulk methylation criterion - which was the lowest level of bulk methylation identifiable using bisulfite (BS)-Seq (8) (see *Materials and Methods*) - and uncovered 14,558 seed DMVs, or 112 Mb of the seed genomic regions, which effectively had no detectable cytosine methylation over all of the developmental stages examined – (Table 1, *Appendix Fig. S1B*, and Dataset S1).

**Table 1.** Summary of Soybean and *Arabidopsis* DMV Characteristics

|  | Soybean |  | *Arabidopsis* |  |
| --- | --- | --- | --- | --- |
|  | < 5%* | < 0.4%* | < 5%* | < 0.4%* |
| Number of DMVs | 21,669 | 14,558 | 4,829 | 3,386 |
| DMV Genome Size | 210 Mb | 112 Mb | 49 Mb | 27 Mb |
| Average DMV Length | 10 ± 6 kb | 8 ± 3 kb | 10 ± 6 kb | 8 ± 4 kb |
| Maximum DMV Length | 76 kb | 41 kb | 137 kb | 39 kb |
| Number of DMV Genes | 14,328 | 6,377 | 8,710 | 3,736 |
| Number of DMV TF Genes | 1,721 | 924 | 835 | 457 |

\* Maximum DMV bulk methylation level in CG-, CHG-, and CHH-contexts across all developmental stages.
† Mean length +/- standard deviation.

**Fig. 1.**
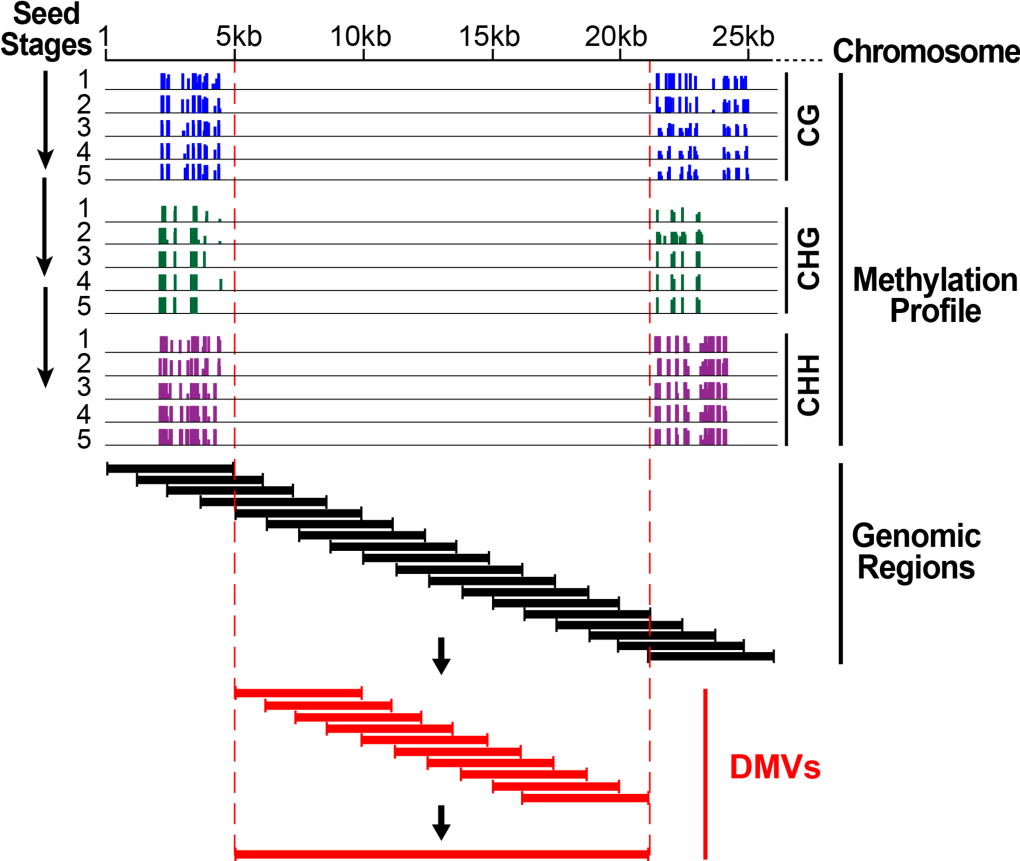
A strategy to identify seed DNA Methylation Valleys (DMVs). Seed methylomes (8) were scanned across the genome at each stage of development (arrows) using a 5 kb sliding window with smaller 1 kb incremental steps (see *Materials and Methods*) (dark bracketed lines). The bulk methylation levels in CG-, CHH-, and CHG-contexts were calculated for each window at every developmental stage (8). Genomic regions with bulk methylation levels of < 5% or < 0.4% across all developmental stages were designated as DMVs, and overlapping DMVs were merged to define DMV regions (red line).

**Fig. 2.**
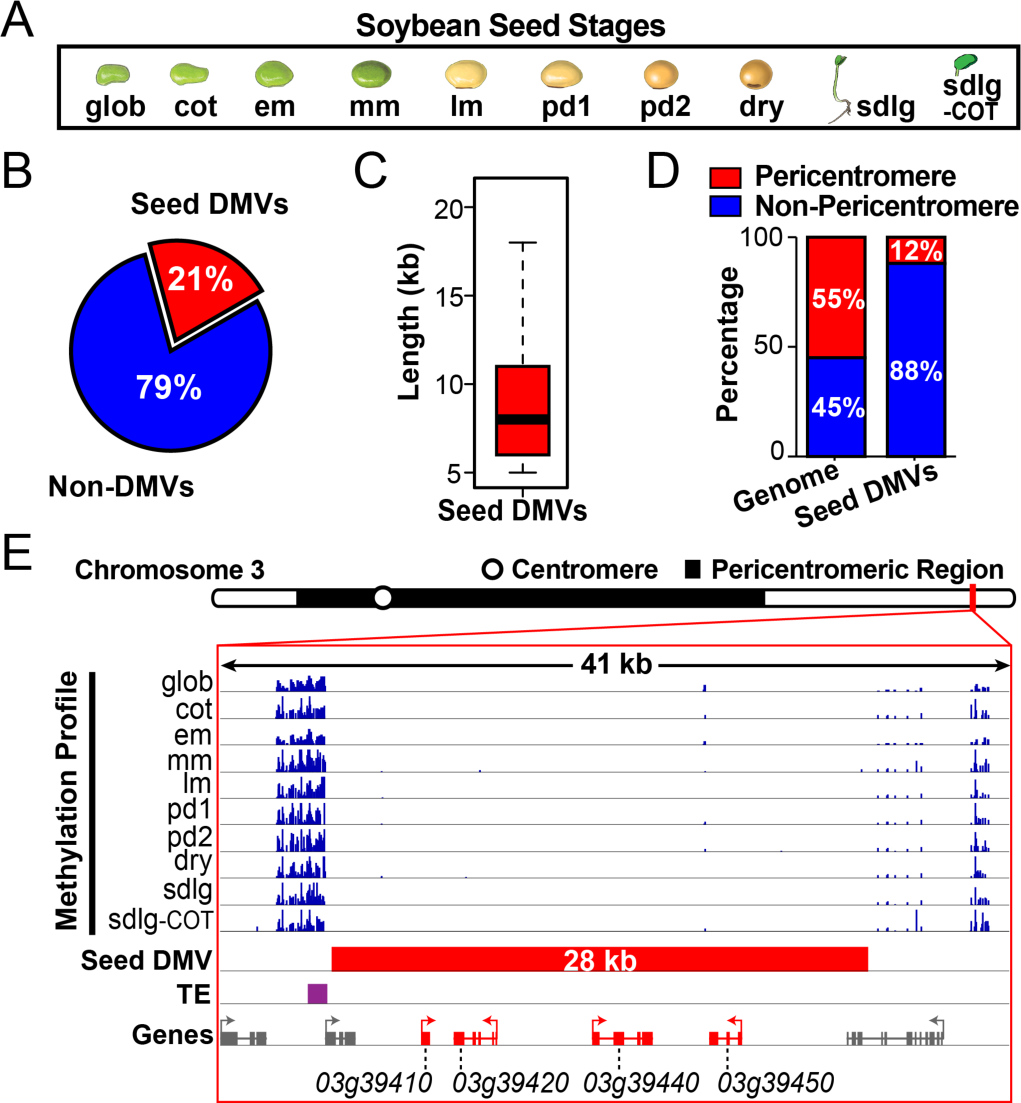
Identification of soybean seed DMVs with < 5% bulk methylation level. (*A*) Seed and post-germination methylomes used to identify DMVs (8). glob, cot, em, mm, lm, pd1, and pd2 refer to globular, cotyledon, early-maturation, mid-maturation, late-maturation, early predormancy, and late pre-dormancy seed developmental stages, respectively. sdlg and sdlg-COT refer to 6 day post-germination seedling and seedling cotyledon, respectively. Seed images are not drawn to scale. (*B*) Percentage of seed DMVs with < 5% bulk methylation in the soybean genome (see *Materials and Methods*). (*C*) Box plot of seed DMV lengths. Horizontal bar represents a median length of 8 kb. (*D*) Percentages of seed DMVs in chromosomal pericentromeric and arm regions. (*E*) Genome browser view of a 28 kb DMV located on chromosome 3. Genes in red color (Glyma03g39410, *EPOXIDE HYDROLASE*; Glyma03g39420, *60S RIBOSOMAL PROTEINL18-3;* Glyma03g39440, *SERINE/THREONINE PROTEIN PHOSPHATASE 2A;* Glyma03g39450, *DORMANCY/AUXIN ASSOCIATED PROTEIN)* are located within this DMV, including 1 kb of 5’ and 3’ flanking regions. Genes in gray color are either located partially or outside this DMV region.

Soybean seed DMVs averaged 10 kb in length, extended up to 76 kb, and 50% of the DMVs ranged in size from 6 to 11 kb at the < 5% scanning criterion (Table 1 and Fig 2C). These values were somewhat smaller using the more stringent < 0.4% scanning criterion (Table 1). Soybean seed DMVs were located primarily on the arms (Fig. *2D)* of all chromosomes (*Appendix Fig. S2*), although over 2,000 DMVs were embedded within highly methylated pericentromeric regions (Fig. 2D). One example of a 28 kb seed DMV that was located on the distal arm of chromosome 3 is shown in Fig. 2*E*. This DMV region had four genes, an average bulk cytosine methylation level of 0.04%, and was static with respect to methylation level during seed development - from the globular stage through dormancy and early germination (Fig. 2*A* and *E*). Taken together, these results indicate that DMVs represent a significant portion of the soybean seed genome, and do not vary significantly with respect to methylation status during seed development and early germination.

**Soybean DMVs Are Present in Different Seed Regions and Tissues**. Soybean DMVs were identified using methylome data from whole seeds (see *Materials and Methods* and Fig. 2A) (8). To determine whether the seed DMVs were present within different seed regions and tissues, we scanned soybean seed methylomes from (i) embryonic axis and cotyledon regions at three developmental stages, and (ii) specific seed coat and cotyledon tissues at early maturation for DMVs at the < 5% methylation criterion (Dataset S1), and compared these DMVs with those identified in seeds as a whole (Fig. 3 *A-C*). We generated these methylomes previously from hand-dissected seed regions (seed coat, axis, and cotyledons), and from specific seed tissues that were captured using laser capture microdissection (LCM) (8).

**Fig. 3.**
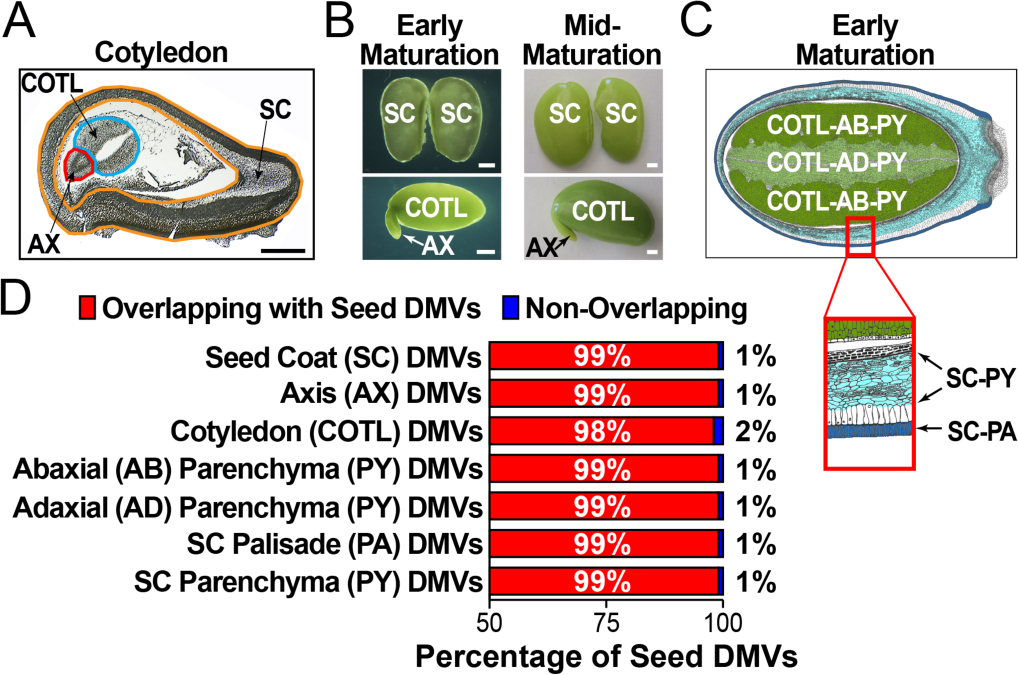
Soybean seed DMVs are present in specific seed regions and tissues. (*A*) Paraffin section of a cotyledon-stage seed. AX, COTL, and SC refer to axis, cotyledon, and seed coat, respectively. *(B)* Whole-mount photographs of embryo and SC from early-maturation and mid-maturation stage seeds. (*C*) Hand drawn colorized cross-section of an early-maturation stage seed. The expanded red box represents an enlarged SC region. COTL-AB-PY, COTL-AD-PY, SC-PA, and SC-PY refer to cotyledon abaxial parenchyma (dark green), cotyledon adaxial parenchyma (light green), seed coat parenchyma (light blue), and seed coat palisade tissues (darkblue), respectively. (*D*) Percentage of seed DMVs that overlap with specific seed region and tissue DMVs. Scale bars indicate 100 um (*A*) and 1 mm (*B*).

The vast majority of whole seed DMVs overlapped with those identified within different seed regions and tissues (Fig. 3*D*). For example, 99% of the whole seed DMVs were found within the seed coat and axis, while 98% were present with the cotyledons. Similarly, 99% of the whole seed DMVs were represented in specific seed coat tissue layers (parenchyma and palisade) and within adaxial and abaxial cotyledon parenchyma tissues (Fig. 3*D*). These results indicate that soybean seed DMVs are not confined to one seed compartment, but are present throughout the seed within diverse regions and tissues that have unique functions and developmental fates, and are a unique feature of seed and seedling genomes.

**Soybean Seed DMVs Are Enriched in Transcription Factor Genes**. We searched seed DMVs for genes, and scored identified genes as DMV genes if they had their gene bodies and 1 kb of 5’ and 3’ flanking regions within DMV regions (see *Material and Methods*, Fig. 2*E*). We uncovered 14,328 and 6,377 diverse genes within DMVs at the < 5% and < 0.4% scanning criteria, respectively (Table 1 and Dataset S2). The former represented 26% of all genes within the soybean genome (Fig. 4*A*).

**Fig. 4.**
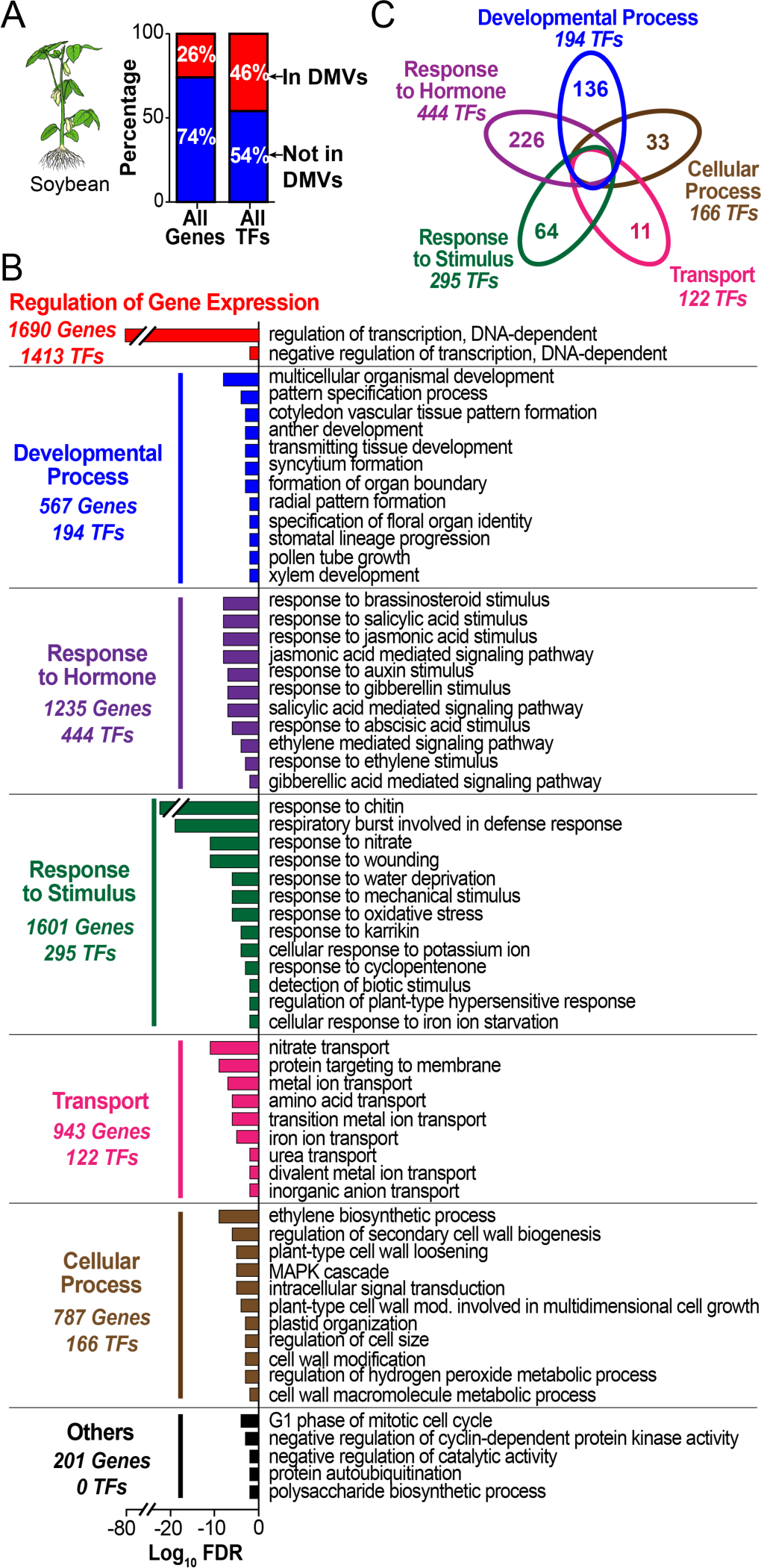
Biological process gene ontology (GO) terms that are enriched in soybean seed DMV genes. *(A)* Proportion of seed DMV genes and transcription factor (TF) genes with < 5% bulk methylation in the soybean genome. *(B)* Enriched GO terms with a Log10 False Discovery Rate (FDR) < 0.05, which are also listed in Dataset S3 (see *Materials and Methods). (C)* Venn diagram showing the number of DMV TF genes that are unique to each of the five GO term biological function groups.

Remarkably, 46% of soybean transcription factor (TF) genes (1,721) were located within seed DMV regions (Fig. 4*A* and Table 1) - a significant enrichment of TF genes within the soybean genome (t-test, P < 0.05). Gene ontology (GO) analysis indicated that the most enriched seed DMV gene GO functional group was regulation of transcription [False Discovery Rate (FDR) 4.9 × 10^−80^], containing 1,413 TF genes, and was consistent with the large number of TF genes represented within seed DMVs (Fig. 4*A*, Table 1, and Dataset S3). A control group of 14,328 randomly selected soybean genes had no GO enrichment terms. DMV genes were distributed into many developmentally and physiologically relevant GO functional classes, such as developmental processes, response to hormones, response to stimulus, and transport, among others (Fig. 4*B*). Seed relevant developmental categories included pattern specification, cotyledon vascular tissue formation, radial pattern formation, organ boundary formation, and multicellular development (Fig. 4*B*). Each of these functional categories contained large numbers of specific TF genes that most likely guide and control these processes (Fig. 4*C*). Together, these data show that TF genes that play major roles in soybean seed formation are preferentially located with DMV regions.

**Many Soybean DMV Genes Are Regulated During Seed Development.** We analyzed soybean whole seed RNA-Seq data, and RNA-Seq data from specific seed regions, subregions, and tissues captured using LCM, for DMV genes that were regulated during seed development (see *Materials and Methods*). We found a large number of DMV genes that were up-regulated > 5-fold in specific seed developmental stages, including many TF genes (Fig. 5*A* and *B*, Dataset S2). For example, *GmWOX9a, GmSPCH*, and *GmTOM3* TF genes were expressed specifically at the globular, early-maturation, and late-maturation stages of seed development, respectively, and were present in DMV regions that did not vary significantly with respect to methylation during seed formation (Fig. 5*C*). All of the DMV stage-specific TF genes (Fig 5*B*) were represented within the regulation of transcription and developmentally relevant functional GO groups (Fig. 4*B* and *C*).

**Fig. 5.**
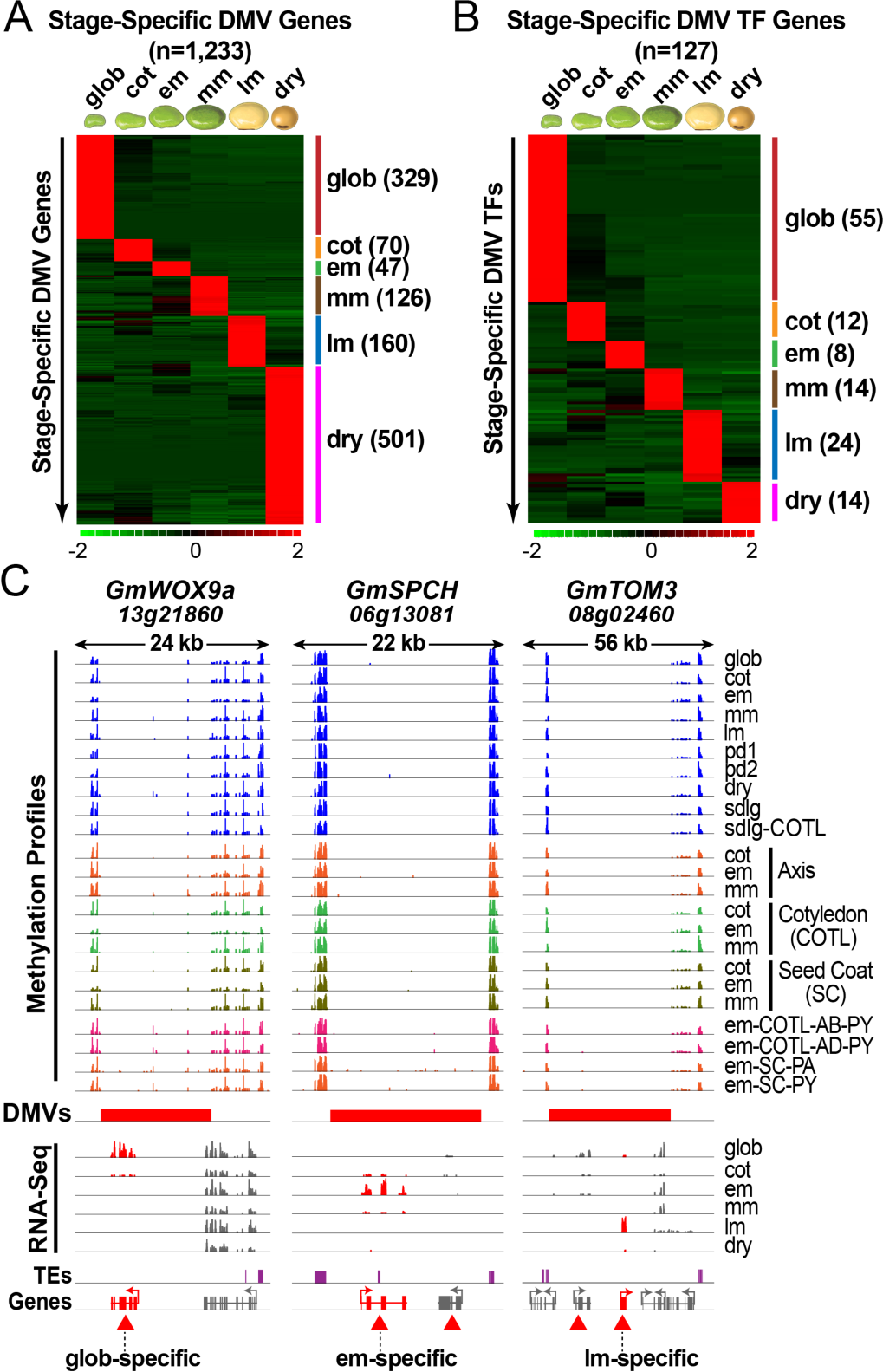
Expression profiles of soybean DMV genes that are up-regulated > 5-fold during seed development. Heat maps of up-regulated seed DMV genes (*A*) and TF genes (*B*) generated by edgeR analysis (33) of soybean RNA-seq whole seed datasets (GSE29163) (8, 23) (see *Materials and Methods*). The number of up-regulated DMV genes and TF genes in each stage is listed on the right side of the heat maps and listed in Dataset S2. Seed images are not drawn to scale. *(C)* Methylome and RNA-Seq genome browser views of three > 5-fold up-regulated stage-specific TF genes *[GmWOX9a, WUSCHEL RELATED HOMEOBOX 9A; GmSPCH, SPEECHLESS; and GmTOM3, TARGET OF MONOPTEROS 3* (red gene models)]. Red triangles mark all genes within the DMVs, including those that are not seed stage-specific. Seed stage, region, and tissue abbreviations are defined in the legend to Figs. 2 and 3. TEs, transposable elements.

A large number of DMV genes, including those encoding TFs, were regulated with respect to specific seed regions, sub-regions, and tissues at different developmental stages as well *(Appendix Fig. S3 A-C* and Dataset S2). Approximately, 22% of all DMV genes (3,189) were expressed preferentially within specific seed parts using a > five-fold up-regulation criterion (*Appendix Fig S3A*) - representing almost half of all soybean seed-region-, subregion-, and tissue-specific genes (*Appendix Fig. S3B*). For example, a *TRIHELIX DNA BINDING PROTEIN* gene located within a 10 kb seed DMV region that was shared by all seed developmental stages, regions, and tissues investigated (8), was expressed specifically within early-maturation stage seed coat palisade tissue layer (*Appendix Fig. S3D and* E). This DMV had a maximum of three methylated cytosines over its entire 10 kb length, did not change its methylation status during development and, was flanked on either side by cliffs of highly methylated transposable elements (*Appendix Fig. S3E*). Taken together, these data show that a large number of soybean genes that reside within seed DMV regions are regulated with respect to space and time and participate in important seed developmental processes (Fig. 4).

**DMV Genes Are Enriched in H27Kme3 and Bivalent Histone Marks**. We investigated soybean embryos at different developmental stages to determine whether DMV genes were coated with specific histone marks (see *Material and Methods*). We used embryos instead of whole seeds to minimize the representation of different cell types in our chromatin preparations. Approximately 90% of all cells were derived from cotyledon parenchyma tissue at all stages of embryo development investigated for histone marks (16).

DMV genes were enriched significantly (hypergeometric test, P < 0.001) in H3K27me3 and bivalent (H3K27me3 and H3K4me3) marks at the cotyledon, early-maturation, mid-maturation, and late-maturation stages, as well as in seedlings, compared with all soybean genes (Fig. 6*A*) - a distinctive feature of animal cell DMVs (12, 13). By contrast, there was no significant enrichment of H3K4me3 at any developmental stage, although a large fraction of expressed DMV genes were coated with this histone mark (Fig. 6*A*). DMV genes marked with H3K4me3 had the highest expression levels, followed by bivalent and H3K27me3 marked genes, respectively (*Appendix Fig. S4*). H3K27me3 marked DMV genes were repressed, or expressed at very low levels, consistent with the repressive nature of the H3K27me3 mark (13). Surprisingly, DMV genes coated with H3K27me3 and bivalent marks were enriched significantly for the transcriptional regulation functional GO term group, a property also similar to their animal DMV gene counterparts (12, 13). For example, at mid-maturation H3K27me3 and bivalent marked DMV genes had FDRs of 2.93 × 10^−24^ and 3.92 × 10^−108^ for this GO term group, respectively. Supporting these results, 62 to 74% of the 1,721 DMV TF genes (Table 1) contained bivalent or H3K27me3 marks depending upon the developmental stage - a significant enrichment compared with all TF genes (hypergeometric test, P < 0.001) (Fig. 6*B*). By contrast, DMV genes marked with H3K4me3 did not generate a transcriptional regulation enrichment group by GO analysis.

**Fig. 6.**
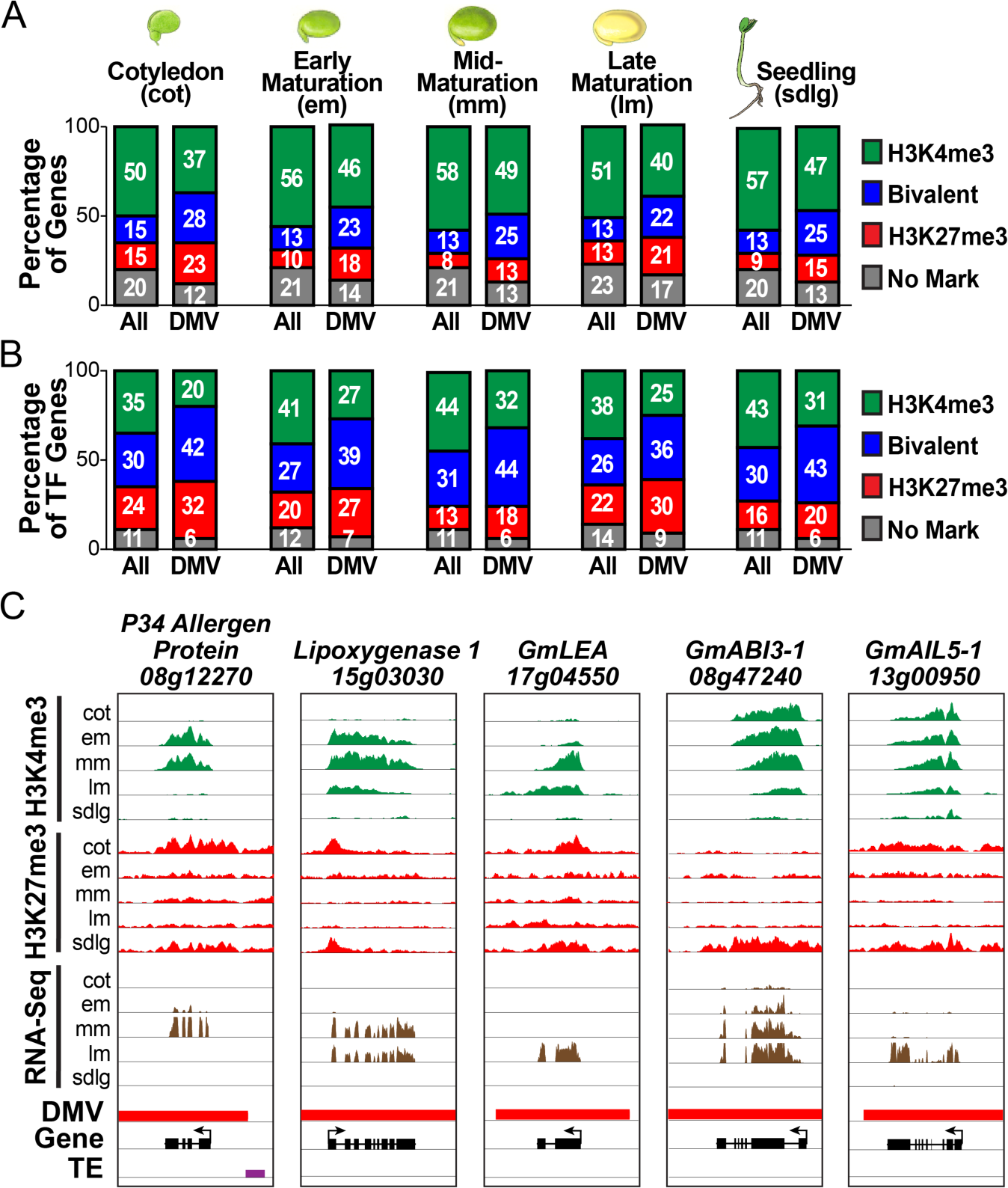
DMV genes with histone marks across soybean seed development. (*A*) Percentage of DMV genes and all genes in the genome marked with H3K4me3, H3K27me3, H3K4me3 and H3K27me3 (bivalent mark), or no mark at each developmental stage (GenBank Accession GSE114879). (*B*) Percentage of DMV TF genes and all TF genes in the genome marked with H3K4me3, H3K27me3, bivalent mark, or no mark at each developmental stage *(C)* Genome browser views showing histone marks and RNA-Seq levels for specific DMV genes during seed development and germination. *GmLEA (LATE EMBRYO ABUNDANT), GmABI3 (ABSCISIC ACID INHIBITOR3), GmAIL5-1 (AINTEGUMENTA LIKE5-1).*

The chromatin marks for many seed DMV genes changed during development in parallel with their expression levels at specific stages (Fig. 6*A* and *Appendix Fig. S4*). For example, the *P34 ALLERGEN PROTEIN* gene had an H3K27me3 repressive mark at the cotyledon stage where it was repressed, an H3K4me3 active mark during early and mid maturation where it was active at a high level, and an H3K27me3 mark in late maturation and post-germination seedlings where it was silent (Fig. 6*C*). The *ABSCISICINHIBITOR3-1 (ABI3-1)* TF gene had an active H3K4me3 mark during seed development when it was expressed, but an H3K27me3 mark following germination when it was repressed (Fig. 6*C*). Finally, the *AINTEGUMENTA-LIKE5-1 (AIL5-1)* TF gene had bivalent marks at cotyledon and early-maturation stages where it was silent or expressed at low levels, an H3K4me3 mark during mid-maturation and late maturation stages where it became active, and a bivalent mark within post-germination seedlings where it was repressed (Fig. 6*C*). Together, these results indicate that (i) seed DMVs are enriched in TF genes that are preferentially coated with H3K27me3 and bivalent marks, (ii) genes within DMV regions undergo modifications in chromatin epigenetic state in the absence of DNA methylation changes during development, and (iii) seed DMVs have features strikingly similar to their animal cell counterparts.

***Arabidopsis* Seeds Also Contain DMV Regions Enriched With Transcription Factor Genes**. We scanned *Arabidopsis* methylomes from seeds at different developmental stages (globular stage through dry seed) and leaves from post-germination plants to determine whether DMVs were a conserved feature of plant genomes (Fig. 7*A*). We used the same strategy to identify *Arabidopsis* seed genomic regions with < 5% and < 0.4% average bulk methylation levels as we did to uncover soybean seed DMVs (see *Materials and Methods*) (Fig. 1). Ninety-nine percent of the DMVs identified during seed development were also shared with leaf DMVs, and, as such, we refer to all DMVs as seed DMVs (see *Materials and Methods*).

**Fig. 7.**
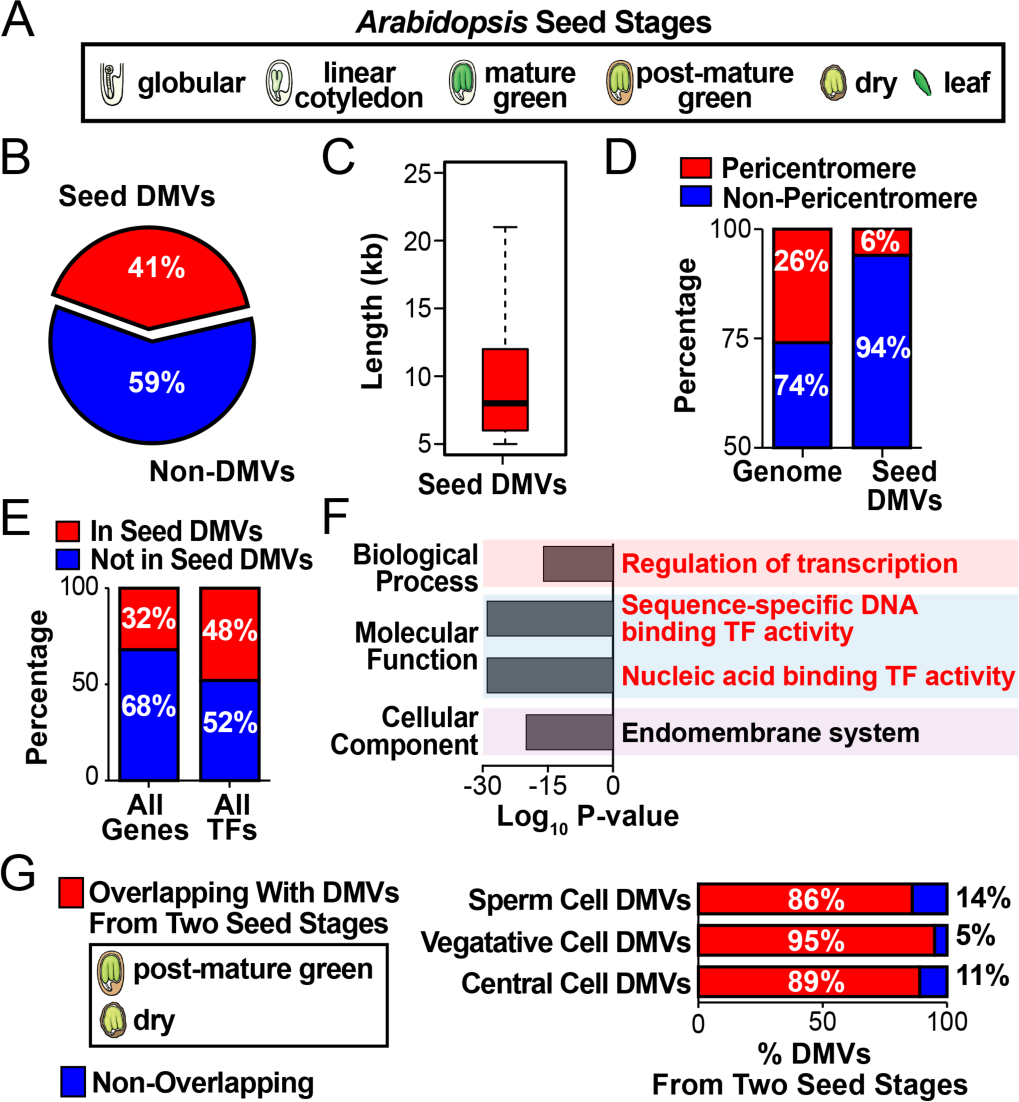
Identification of *Arabidopsis* seed DMVs with < 5% bulk methylation level. *(A) Arabidopsis* methylomes used to identify DMVs (8). Images are not drawn to scale. *(B)* Proportion of seed DMVs with < 5% bulk methylation level in the *Arabidopsis* genome. *(C)* Box plot of seed DMV lengths. Horizontal bar represents a median length of 8 kb. *(D)* Proportion of seed DMVs in chromosomal pericentromeric and arm regions. *(E)* Proportion of seed DMV genes and DMV TF genes with < 5% bulk methylation in the *Arabidopsis* genome. *(F)* Bar plots of the most significantly enriched GO terms of *Arabidopsis* seed DMV genes, which are also listed in Dataset S4. Log10 P-value refers to FDR. GO terms related to transcription factor activity are highlighted in red. (*G*) Percentages of seed DMVs that overlap with sperm cell, vegetative cell, and central cell DMVs.

Approximately 41% of the *Arabidopsis* genome, or 4,829 regions, consisted of seed DMVs at the < 5% scanning criterion (Fig. 7*B*, Table 1, and Dataset S1). These regions had a bulk cytosine methylation level of 0.24%, on average, compared with 5.8% for the *Arabidopsis* genome as a whole *(Appendix Fig. S1C and D*). *Arabidopsis* DMV regions averaged 10 kb in length and were localized primarily on the arms of all chromosomes, similar to what was observed in soybean (Table 1, Figs. 7*C* and *D*). A smaller number of seed DMVs (3,386) were uncovered using the < 0.4% scanning criterion that identified genomic regions with essentially no methylation (Table 1).

Approximately 32% of all *Arabidopsis* genes (8,710), including 48% of those encoding TFs (835), were localized within seed DMV regions at the < 5% scanning criterion (Fig. 7*E*, Table 1, and Dataset *S2). Arabidopsis* TF genes were enriched significantly within DMVs (t-test, P < 0.05), and about one-half overlapped with those present within soybean DMV regions. GO enrichment analysis of all *Arabidopsis* seed DMV genes showed that the most significant GO functional groups were sequence-specific DNA binding TF activity, nucleic acid TF binding activity, and regulation of transcription (Fig. 7*F* and Dataset S4), similar to what was observed for soybean DMVs (Fig. 4). Other developmentally relevant GO functional groups, such as response to hormones, also showed significant enrichment (*Appendix Fig. S5* and Dataset S4). DMV genes shared between *Arabidopsis* and soybean showed GO enrichment for many biological processes related to development, as predicted from the DMV GO analysis of each plant species separately *(Appendix Fig. S6).* Finally, large numbers of *Arabidopsis* genes that reside within DMVs, including those encoding TFs, were regulated with respect to time and space during seed development *(Appendix Figs. S7 and S8*, and Dataset S2). Together, these results show that *Arabidopsis* seed DMV regions have features identical to those in soybean seeds, and that DMVs enriched for TF genes and other genes that carry out important seed developmental functions are a conserved feature of seed genomes.

**Most Seed DMV Regions Are Present Before Fertilization**. In both soybean and *Arabidopsis*, 99% of DMVs uncovered during seed development were conserved within the genomes of seedlings (soybean) and leaves *(Arabidopsis)*, indicating that the seed DMV methylation status did not change significantly following germination (see *Materials and Methods*). We scanned *Arabidopsis* sperm, vegetative, and central cell methylomes generated by others for DMVs (17, 18) (Fig. 1), and compared DMVs present in these gametophytic cells with those uncovered from our *Arabidopsis* post-mature green and dry seed methylomes (8) to determine whether seed DMVs were present prior to fertilization (Fig. 7*G*). We used seed methylomes from the Col-0 ecotype (Fig. 7*G*) instead of Ws-0 (Fig. *7A-F)*, because the gametophytic cell methylomes were from Col-0, and we found that 10% of Ws-0 seed DMVs differed from those in Col-0 at the same developmental stages due to cytosine polymorphisms (8, 19, 20). Thus, it was important to carry out within ecotype DMV comparisons.

The vast majority (86 to 95%) of *Arabidopsis* seed DMV regions were also present within female gametophyte (central cell) and male gametophyte (sperm and vegetative cells) genomes prior to fertilization (Fig. 7*G*). The small number of seed DMV regions that were not scored as DMVs in sperm and central cells had average bulk methylation levels > 5%, and, therefore, higher methylation levels before fertilization and seed development. These results indicate that most seed DMV regions are highly conserved, and do not change significantly with respect to methylation status during major periods of the plant life cycle. They are present within gametophytic cells before fertilization, developing seeds after fertilization, growing sporophytic seedlings following seed germination, and leaves of the young sporophytic plant.

**Many Genes Known to Play Important Roles in Seed Development Are Present Within Soybean and *Arabidopsis* DMVs**. GO analysis indicated that genes contained within both soybean and *Arabidopsis* DMVs were significantly enriched for those that play major regulatory roles during seed formation (Figs. 4 and 7*F*, *Appendix Figs. S5 and S6*). We searched soybean and *Arabidopsis* DMVs for genes that were known to play critical roles in seed differentiation and development (Dataset S5). Many genes essential for embryo formation were localized within both soybean and *Arabidopsis* DMVs - including several *WUS HOMEOBOX-CONTAINING (WOX)* genes (e.g., *WOX1, WOX3, WOX8/9), BABY BOOM, SCARECROW, SHATTERPROOF1, PLETHORA, TARGET OF MONOPTEROS5*, and *CUP SHAPED COTYLEDON* family genes (e.g., *CUC2, CUC3*). Other genes playing major roles in seed formation, such as *CLAVATA3, PIN1, YUCCA4, BODENLOS*, and *HANABA TARANU*, were also present within both soybean and *Arabidopsis* DMVs. In addition, several gene classes that play critical roles in seed physiological processes, such as those encoding storage proteins (e.g., *GmGlycinin1, AtCruciferin1)* and hormone biosynthesis enzymes [e.g., Gibberellic Acid Oxidase (*GmGA20Ox2, GmGA3Ox3, AtGm20Ox2, AtGA3Ox3*)] were also localized with both soybean and *Arabidopsis* DMVs. Together, these results suggest that many genes playing essential roles in seed formation are conserved within the DMVs of divergent plant species.

## Discussion

We have shown that there are large portions of soybean and *Arabidopsis* seed genomes that are hypomethylated and enriched with TF genes. Seed DMVs resemble in striking ways the DMVs that are present in the cells of many vertebrate animals, including humans (12-14, 21). DMVs in both kingdoms are (i) scattered across the genome in thousands of long hypomethylated regions, (ii) do not vary significantly with respect to their methylation status across diverse developmental stages, tissues, and cell types, (iii) are enriched significantly with developmentally important genes, including large numbers of TF genes, (iv) undergo epigenetic changes at the chromatin level that can accompany changes in gene activity, and (iv) present within diverse species. Thus, DMVs appear to be a unique feature of eukaryotic genomes that play important roles in cell differentiation and development in the absence of methylation changes at the DNA level. It has been proposed that DMVs arose in animal cells as a result of strong negative selection, or purification, for transposable element insertions that would disrupt critical regulatory genes and have a deleterious effect on development (22). Whether this is the case for the seed DMVs uncovered here remains to be determined.

A large number of genes within both soybean and *Arabidopsis* DMVs are regulated during seed development, and are expressed within specific seed stages, regions, subregions, or tissues. These genes include those encoding TFs, as well as others, such as storage protein genes, which play important roles in seed formation and germination. The remarkable feature of these genes is that they retain their hypomethylated status within DMVs regardless of their state of expression. How then are these genes turned on and off in the absence of methylation changes that have been shown to play, for example, a role in differential activity of maternal and paternal alleles which is essential for seed endosperm development (7)?

One possibility is that many DMV genes, especially those encoding storage proteins, are activated and repressed by the action of seed-specific regulators such as *LEAFY COTYLEDON1 (LEC1), ABSCISIC ACID INHIBITOR3 (ABI3)*, and *FUSCA3 (FUS3)* (23), among others. Epigenetic changes at the chromatin level within seeds may also partner with the action of TFs to facilitate development-specific changes in gene activity. Soybean seed DMV genes, like their animal counterparts (12, 13, 15), are enriched for H3K27me3 and bivalent marks, and the histone marks coated on DMV genes, including H3K4me3, can change in parallel with their gene activity levels. DMV genes that are repressed, or expressed at low levels, are coated with repressive H3K27me3 marks, whereas those that are active are marked with either H3K4me3 or bivalent marks. We assume that the bivalent marks on seed DMV genes resemble those that coat animal DMV genes, rather than being caused by a mixture of tissues in which the genes are active and repressed. First, seed DMV genes coated with bivalent marks are enriched with TF genes, similar to their animal counterparts (24). Second, seed DMV genes that contain bivalent marks are expressed at significantly lower levels than their counterparts marked with H3K4me3 - which is a feature of animal genes with bivalent marks (24). Finally, seed DMV genes marked with both H3K27me3 and H3K4me3 were identified from soybean embryos that consist primarily of parenchyma storage tissue cells that reside within cotyledon regions (Fig 3). Thus, we favor a model in which the actions of TFs coupled with chromatin-level epigenetic changes regulate genes within seed DMV regions during development in the absence of DNA methylation events. In animals, TF genes have been shown to regulate other genes within the same DMV, and are organized into self-contained chromatin domains (15). Whether that is also true of seed DMVs remains to be determined.

What about seed genes that are not present within DMVs? These genes reside within genomic regions with > 5% average bulk methylation at CG, CHG, and/or CHH sites. These regions contain highly methylated transposable elements, genes with body methylation, or both (8). Many soybean and *Arabidopsis* genes outside of DMV regions are also regulated with respect to space and time during seed development (8). We showed previously that many of these genes are activated and shut down in the absence of significant methylation changes similar to genes within DMV regions (8). Whether these genes undergo major epigenetic chromatin changes during seed development remains to be determined.

A major challenge for the future will be to determine the precise mechanisms by which genes residing within both DMV and non-DMV regions are regulated during seed development. This will require identifying the cis-control elements that program seed gene activity, the cognate TFs that interact with these elements, proteins that drive epigenetic changes at the chromatin level, and how genes expressed during seed development are organized into regulatory networks that are required to form a seed.

## Materials and Methods

**General Strategy For Identifying DMVs**. Soybean (*Glycine max* cv Williams 82) and *Arabidopsis* (Ws-0 and Col-0) seed and post-germination methylome data were taken from our previously published BS-Seq experiments (8). Specific details regarding the BS-Seq libraries used in the experiments reported here, and how they were constructed and characterized, are contained within the *Materials and Methods* and *SI Materials and Methods* reported in Lin *et al* (8).

DMVs were identified using the strategy illustrated in Fig. 1 (12). The methylome at each developmental stage was scanned using a sliding window of 5 kb with smaller 1 kb incremental steps, and regions without cytosine bases were discarded. The bulk methylation levels in CG-, CHG-, and CHH-contexts were calculated for all remaining windows (8, 25). DMVs were defined as genomic regions with either < 5% or < 0.4% bulk methylation levels in all three cytosine contexts over all developmental stages studied (Table 1). A 0.4% bulk methylation level was used to identify DMVs with no detectable methylation, as this was the lowest level of un-methylated cytosines that were not converted by BS to thymine in our previous experiments (8). Overlapping DMVs were merged and reported as contiguous DMV regions in Dataset S1. Only genes that contained their entire bodies and 1 kb of 5’ and 3’ flanking regions were designated as genes contained within DMV regions (Dataset S2).

**Identification and Characterization of Soybean Seed DMVs**. Soybean DMVs were identified in the methylomes of globular, cotyledon, early-maturation, mid-maturation, late-maturation, early-pre-dormancy, late-pre-dormancy, and dry seeds, as well as 6 d post-germination seedlings and seedling cotyledons (Fig. 2). Ninety-nine percent of the DMVs identified during seed development was also present as DMVs in post-germination seedlings and seedling cotyledons. Thus, we refer collectively to these DMVs as soybean seed DMVs (Fig. 2 and *Dataset S1*).

***GO term enrichment analysis of seed DMV genes**.* The (i) GOSeqR Bioconductor package (26), (ii) SoyBase GO annotations (https://soybase.org/genomeannotation/index.php), (iii) hypergeometric statistical test method, and (iv) a FDR < 0.05 (Benjamini-Hochberg multiple testing correction) were used to identify GO terms enriched in soybean DMV genes (Fig. 4 and Dataset S3) (23).

***Expression analysis of seed DMV genes**.* dChip (27) was used to generate heat maps of soybean up-regulated DMV mRNA levels in whole seeds (Fig. 5) and in specific seed regions, sub-regions, and tissues *(Appendix Fig. S3)* throughout development. Whole seed RNA-Seq data were taken from our previously published experiments (8) (GenBank Accession GSE29163). RNA-Seq data for specific seed regions, sub-regions, and tissues were taken from the Harada-Goldberg LCM datasets (GeneBank Accessions GSE116036) (23). EdgeR was used to identify DMV mRNAs that were up-regulated > 5-fold [FDR < 0.001 (Benjamini-Hochberg multiple testing correction)] in specific seed stages, regions, sub-regions, and tissues (23).

***Identification of histone marks on DMV genes**.* Embryos at different developmental stages were used to characterize DMV genes for H3K4me3 and H3K27me3 histone marks (Fig. 6). ChIP assays were carried out using anti-H3K4me3, anti-H3K27me3, and anti-H3 antibodies according to our previously published protocol (23). ChIP-Seq library construction, DNA sequencing analysis, and peak calls were carried out as described previously (23). H3 ChIP-Seq results were used as a control to filter out noise during peak calling. MACS2 and SICER were used to call H3K4me3 and H3K27me3 peaks, respectively (28-30). DMV genes were considered marked with H3K4me3, H3K27me3, or both H3K4me3 and H3K27me3 (bivalent marks) if peaks had a P value of < 0.05, and were associated with the gene body and/or 1 kb of upstream region (23). All ChIP-Seq data reported in this paper were deposited in GenBank as accession GSE114879.

**Identification and Characterization of *Arabidopsis* Seed DMVs***. Arabidopsis* (Ws-0) DMVs were identified in the methylomes of globular, linear cotyledon, mature green, post-mature green, and dry seeds, as well as leaves from four week-old plants (8) (Fig. 7). Ninety-nine percent of the DMVs identified during seed development was also present as DMVs in leaves. Thus, we refer collectively to these DMVs as *Arabidopsis* seed DMVs (Fig. 7 and Dataset S1). *Arabidopsis* (Col-0) methylomes from post-mature green and dry seeds (8), and those from sperm, vegetative, and central cells (17, 18) were used to compare DMVs from seeds and gametophytic cells.

***GO term enrichment analysis**.* Virtual Plant 1.3 (31) was used for *Arabidopsis* gene GO enrichment analysis with a cutoff FDR value < 0.05 (Benjamini-Hochberg multiple testing correction) (Fig. 7 and *Appendix Fig. S5 and S6*).

***Gene expression analysis**.* dChip (27) was used to generate heat maps of *Arabidopsis* DMV mRNA levels in whole seeds (Fig. S7) and in specific seed regions and subregions (*Appendix Fig. S8)* during development. *Arabidopsis* seed transcriptome data were taken from our previously published GeneChip experiments [whole seeds (GeneBank Accession GSE680) (32); regions and subregions GeneBank Accession GSE12404 (4)].

## ACKNOWLEDGEMENTS

This work was supported by a grant from the National Science Foundation Plant Genome Program (to R.B.G., M.P., and J.J.H.).

---

Supplementary Information for References for SI reference citations **Other supplementary materials for this manuscript include the following:**

**Fig. S1.**
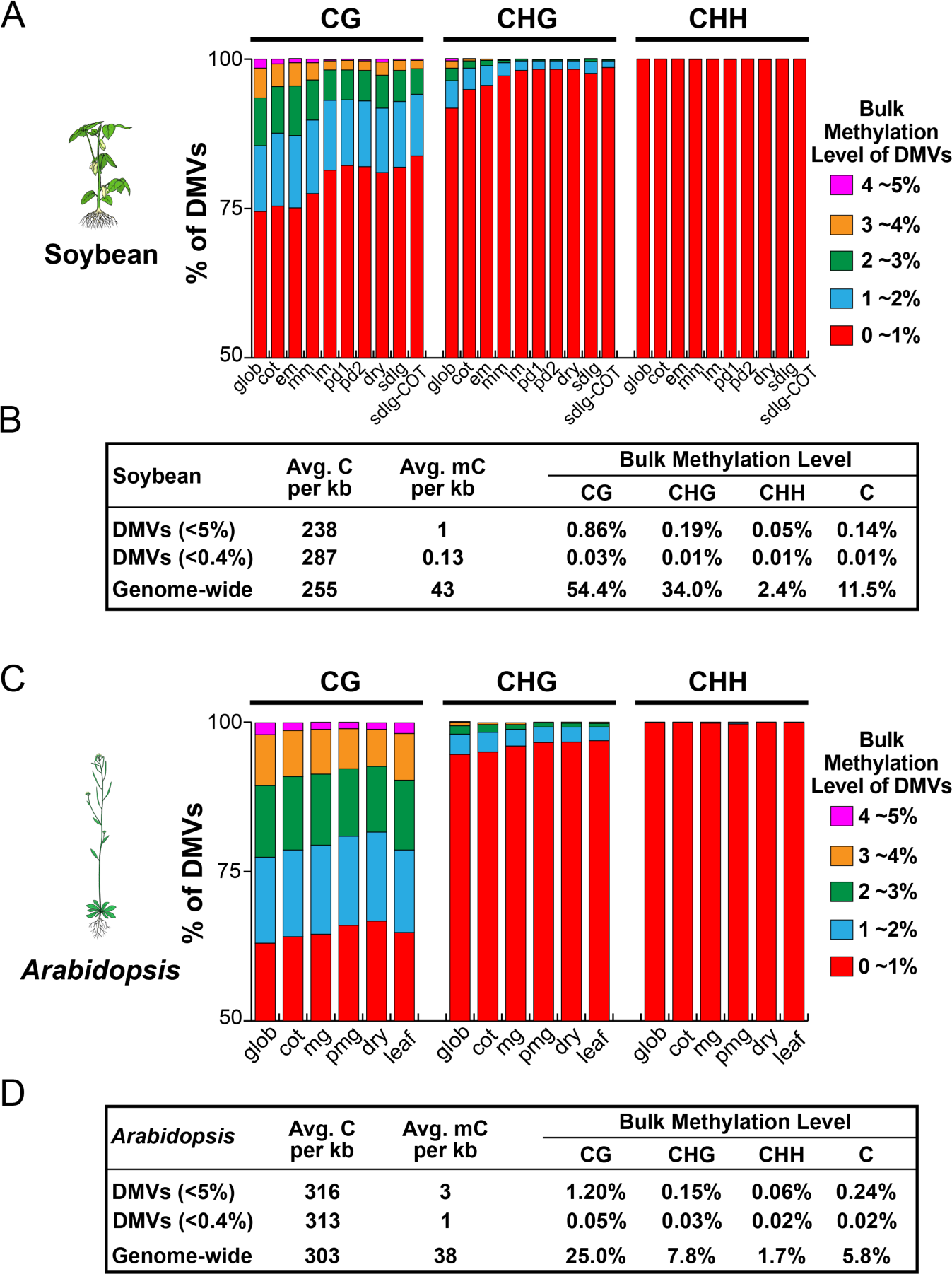
Bulk methylation levels of soybean and *Arabidopsis* DMVs. Average bulk methylation levels were calculated as described previously (see *Materials and Methods*) (1).

**Fig. S2.**
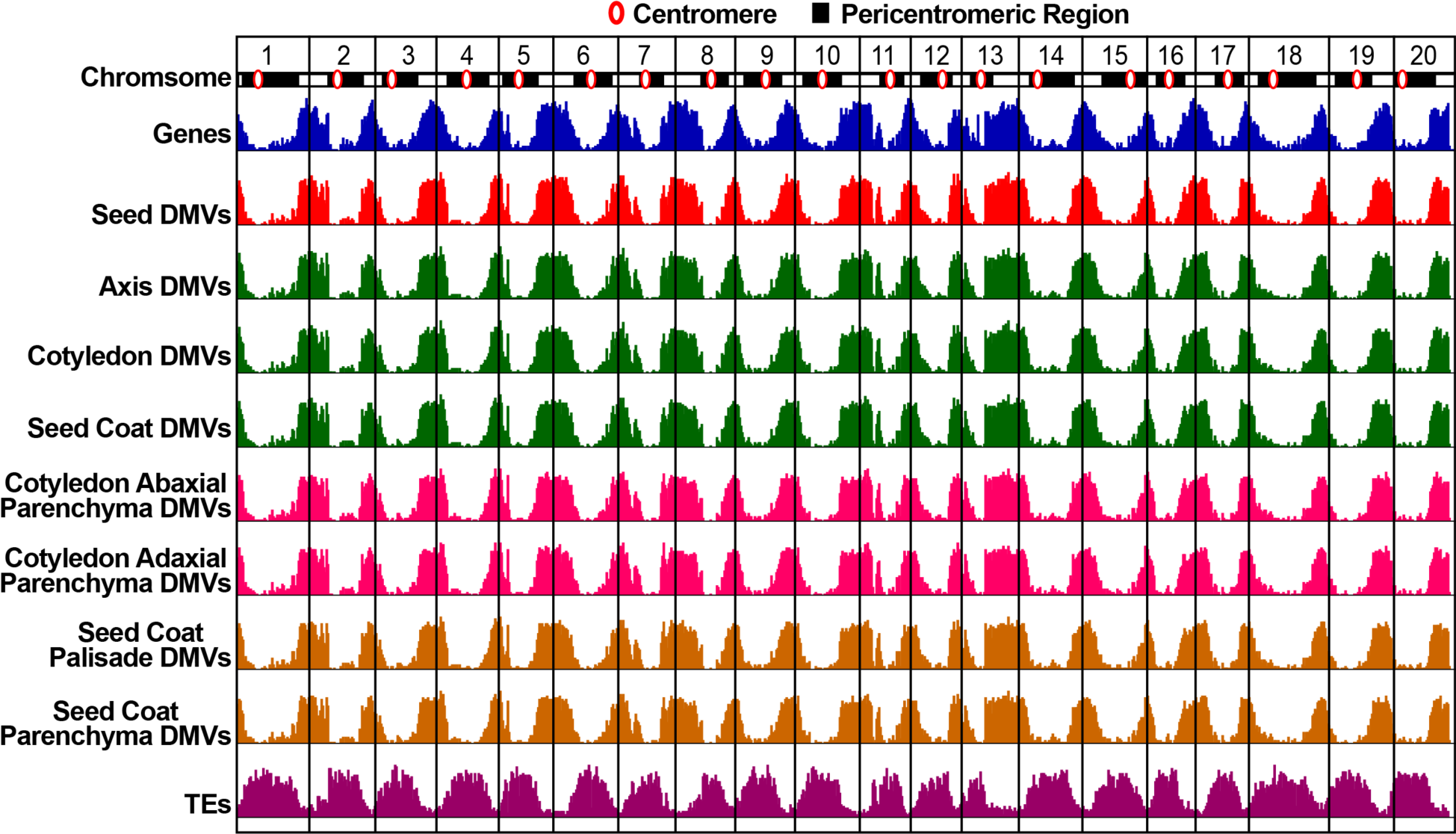
Genome browser view of DMVs, genes, and transposable elements (TEs) across all 20 soybean chromosomes.

**Fig. S3.**
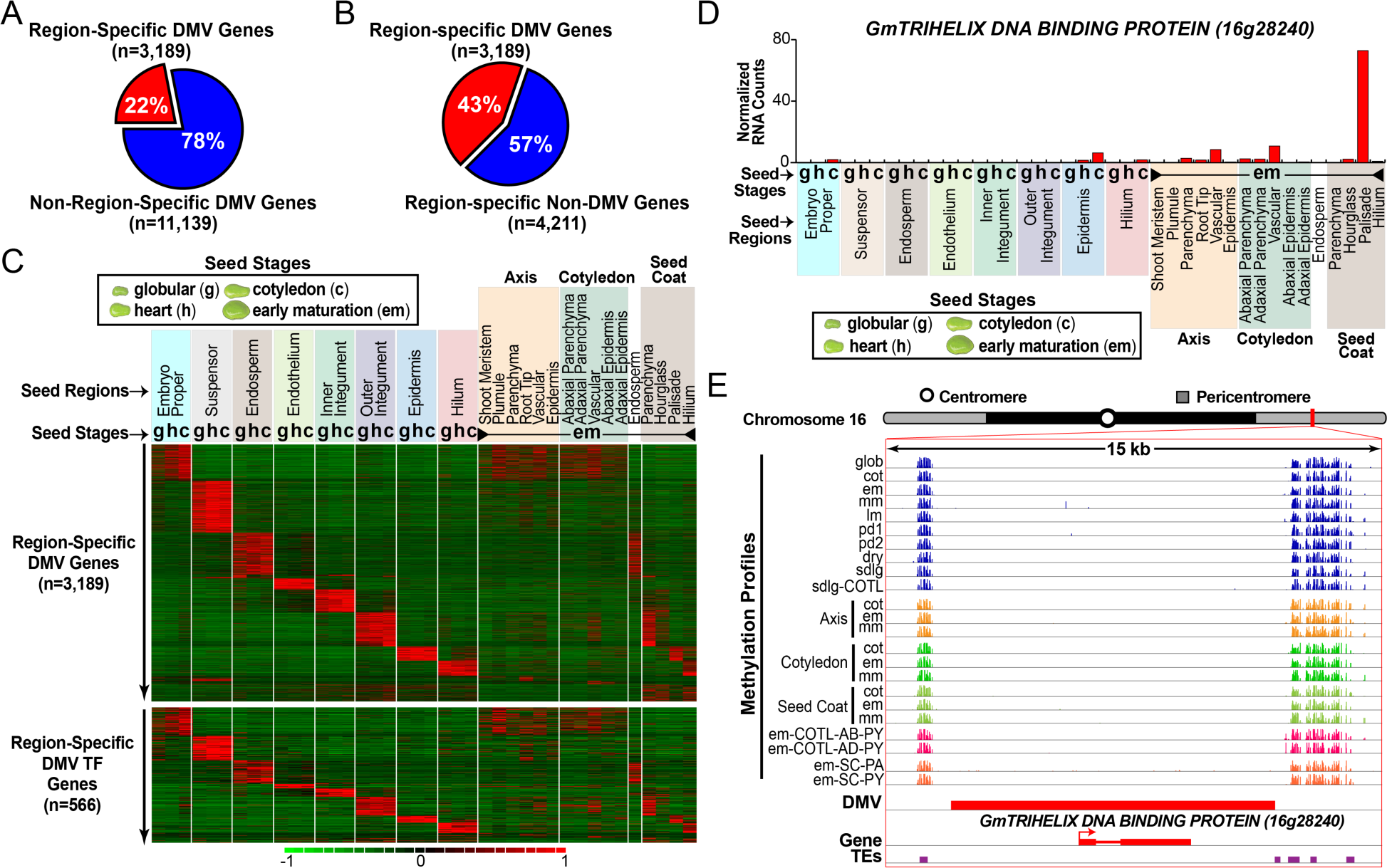
Expression profiles of soybean DMV genes that are up-regulated in specific seed regions, sub-regions, and tissues during development. *(A)* Percentage of DMV genes (14,328) that are up-regulated > 5-fold in specific seed regions, sub-regions, or tissues during development based on edgeR analysis (2) of the Harada-Goldberg LCM soybean seed RNA-seq datasets (GSE116036). (3). *(B)* Percentage of all > 5-fold up-regulated seed region-, subregion-, and tissue-specific genes (7,400) that are in DMVs. *(C)* Heat maps of DMV genes that are up-regulated > 5-fold in specific seed regions, sub-regions, or tissues during development [red sectors in (*A*) and (*B*)]. Expression profile *(D)* of a *TRIHELIX DNA BINDING PROTEIN TF* gene that is specific for the SC palisade layer (Fig. 3C), and the genome browser view *(E)* of its DMV region. The bulk methylation level of this DMV was < 0.01%. Seed images are not drawn to scale. g and glob, globular stage; c and cot, cotyledon stage; h, heart stage; em, early-maturation stage; mm, mid-maturation stage; lm, late-maturation stage; pd1, early predormancy stage; pd2, late pre-dormancy stage; sdlg, seedling; COTL, cotyledon; SC, seed coat; AB, abaxial; AD, adaxial; PY, parenchyma; PA, palisade; TE, transposable element.

**Fig. S4.**
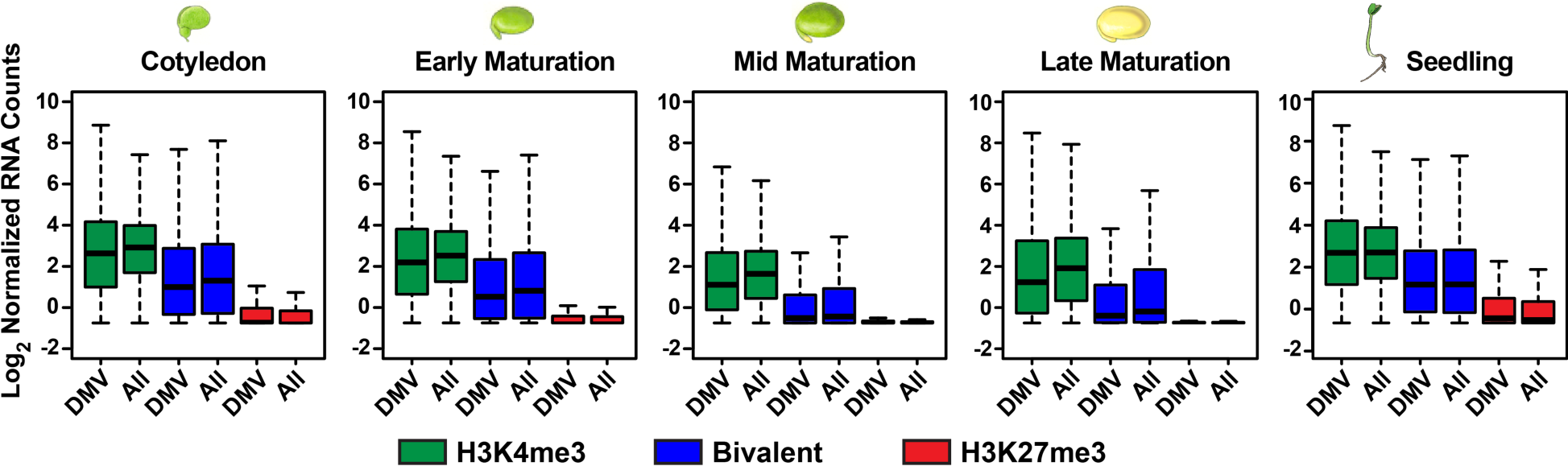
Box plots of RNA-Seq counts for DMV genes and all genes in the soybean genome marked with H3K4me3, H3K27me3, and bivalent marks during seed development. RNA levels for genes with bivalent and H3K27me3 marks differed significantly from the RNA levels of H3K4me3 marked genes (t-test, P < 0.001).

**Fig. S5.**
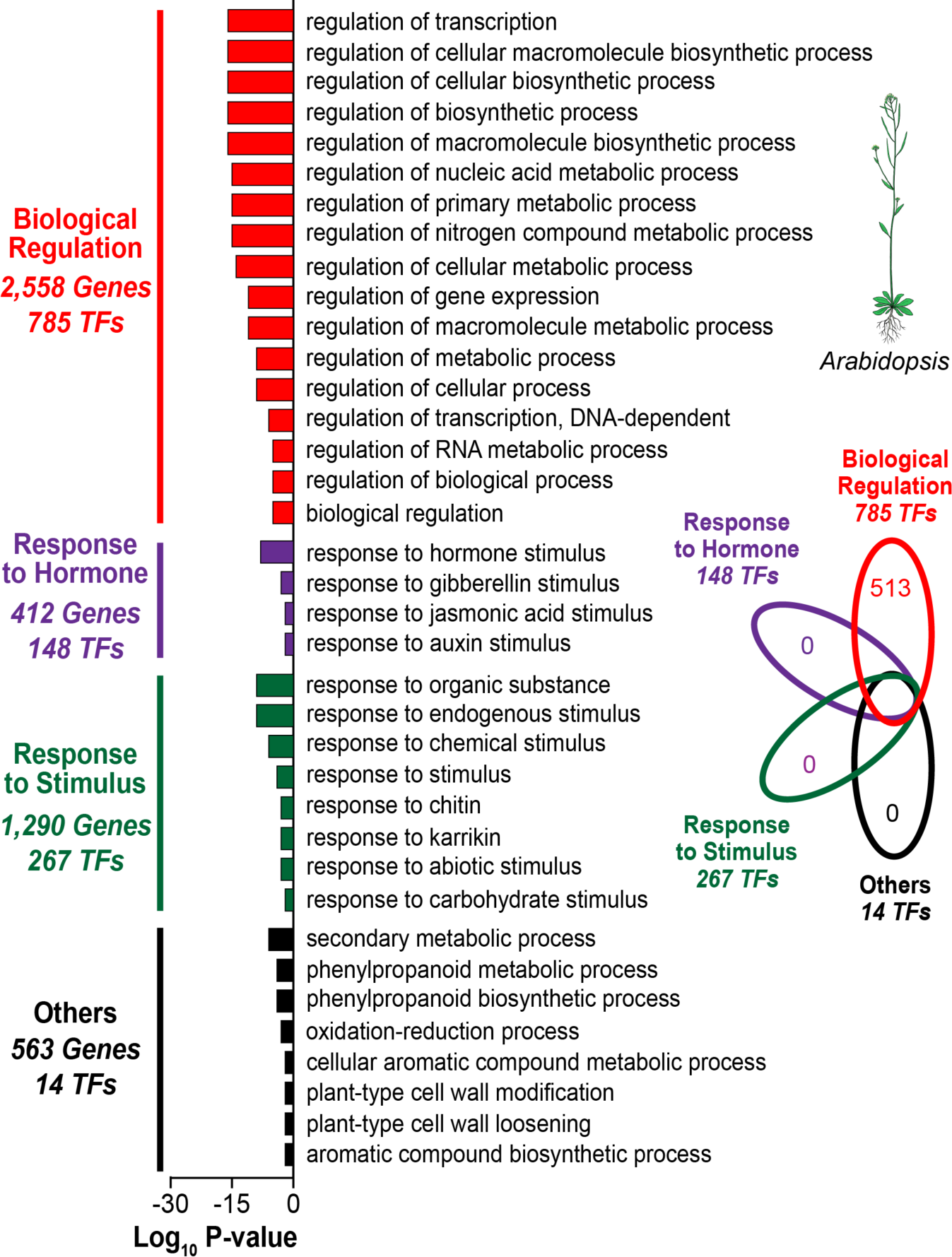
Biological process GO terms that are enriched in *Arabidopsis* seed DMV genes. Enriched GO terms with FDR < 0.05 are also listed in Dataset S4 (see *Materials and Methods*). Venn diagram shows the number of DMV TF genes that are unique to GO term biological function groups.

**Fig. S6.**
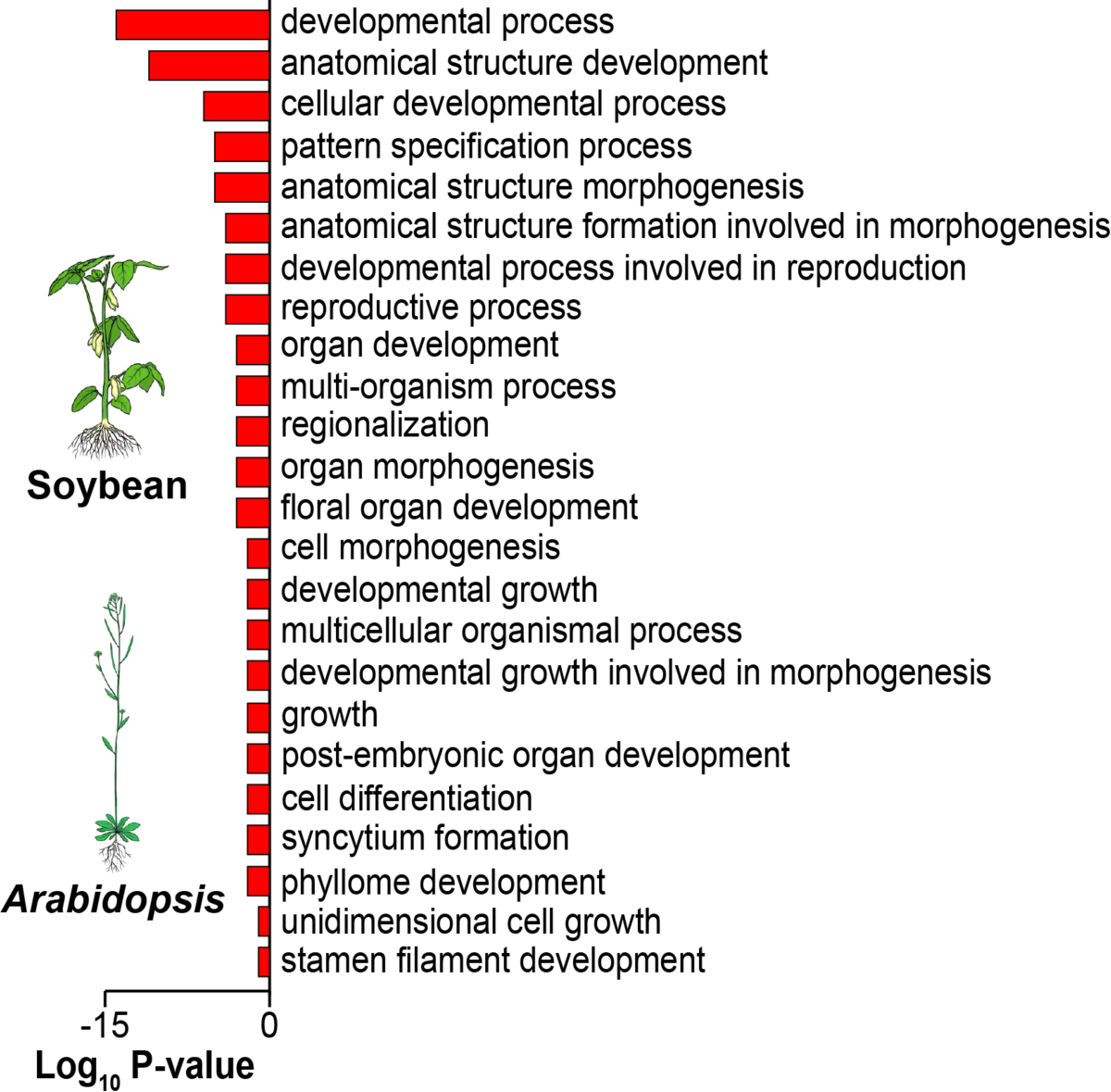
Biological process GO terms related to development that are enriched in seed DMV genes shared between *Arabidopsis* and soybean. *Arabidopsis* gene ID numbers were obtained for DMV genes shared between soybean and *Arabidopsis.* GO enrichment analysis with a cutoff FDR value < 0.05 was carried out as described for *Arabidopsis* genes in *Materials and Methods.*

**Fig. S7.**
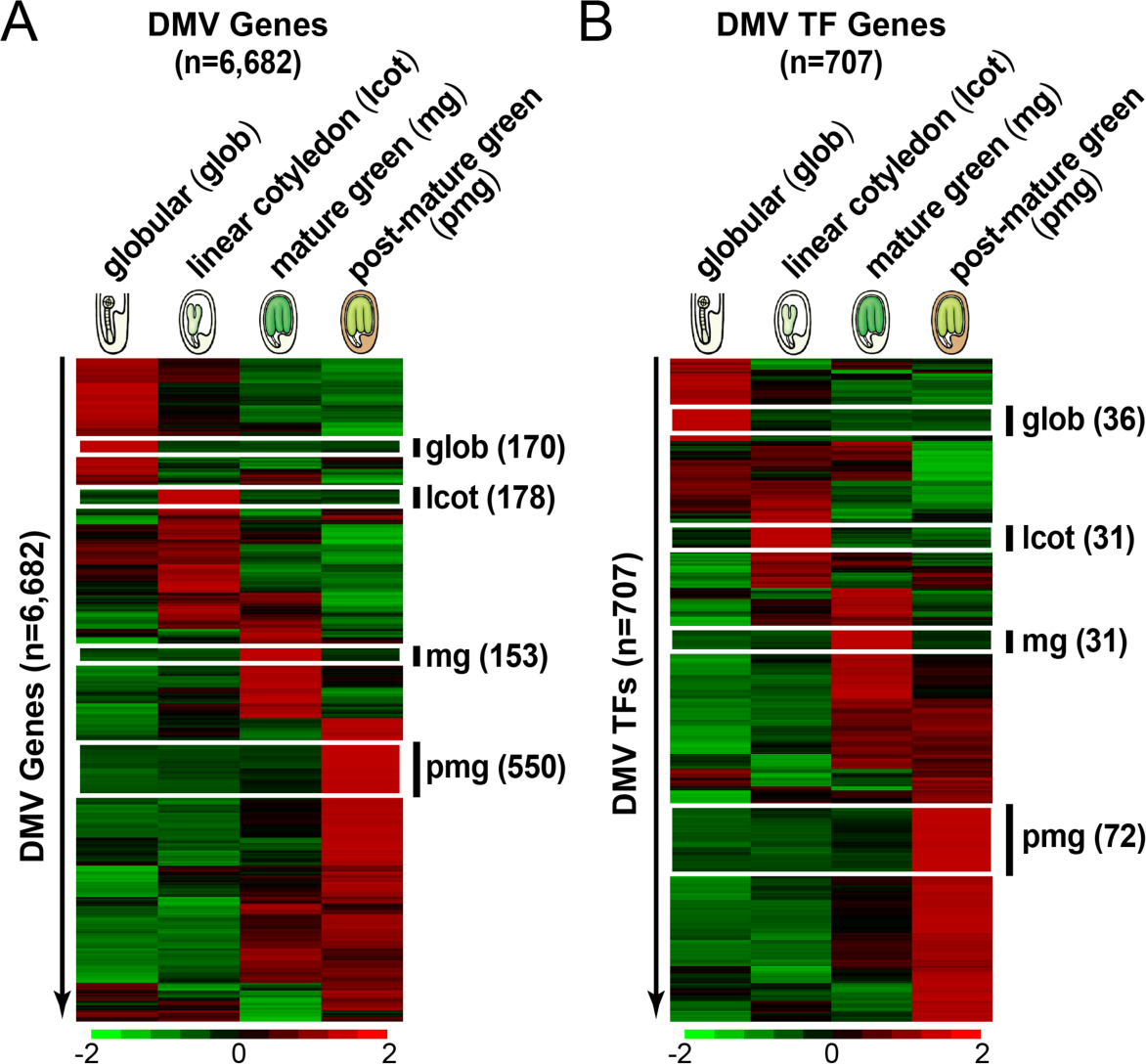
Expression profiles of *Arabidopsis* DMV genes at different stages of seed development. Hierarchical clustering of *Arabidopsis* DMV mRNAs (*A*) and DMV transcription factor mRNAs (TFs) *(B)* during seed development using dChip and our previously published *Arabidopsis* whole seed mRNA GeneChip data (4). Only 6,682 of the 8,710 *Arabidopsis* DMV genes (Table 1) are represented on the ATH1 22k GeneChip used to obtain these expression profiles (4). The number of DMV genes represented in stage-specific RNA clusters is listed on the right side of each figure. Seed images are not drawn to scale.

**Fig. S8.**
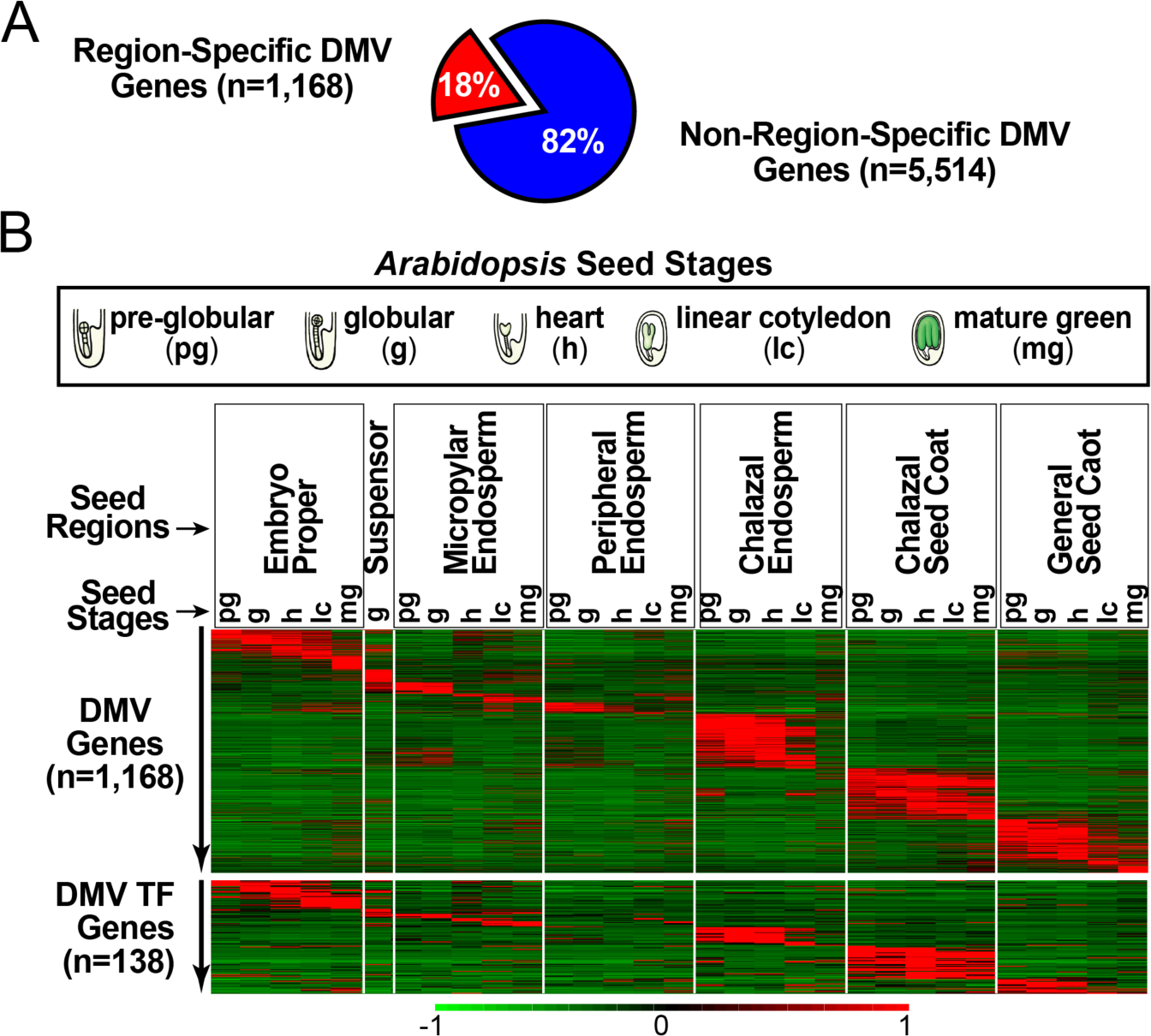
Expression profiles of *Arabidopsis* DMV genes that are up-regulated in specific seed regions and subregions during development. *(A)* Percentage of DMV genes (6,682) that are up-regulated > 5-fold in specific seed regions and subregions at different developmental stages based on our previously published *Arabidopsis* seed LCM mRNA data using the ATH1 22k GeneChip (5). (*B*) Heat maps of DMV mRNAs and those encoding transcription factors (TFs) that are up-regulated > 5-fold in specific seed regions and subregions during development. Seed images are not drawn to scale.

**Dataset S1.**
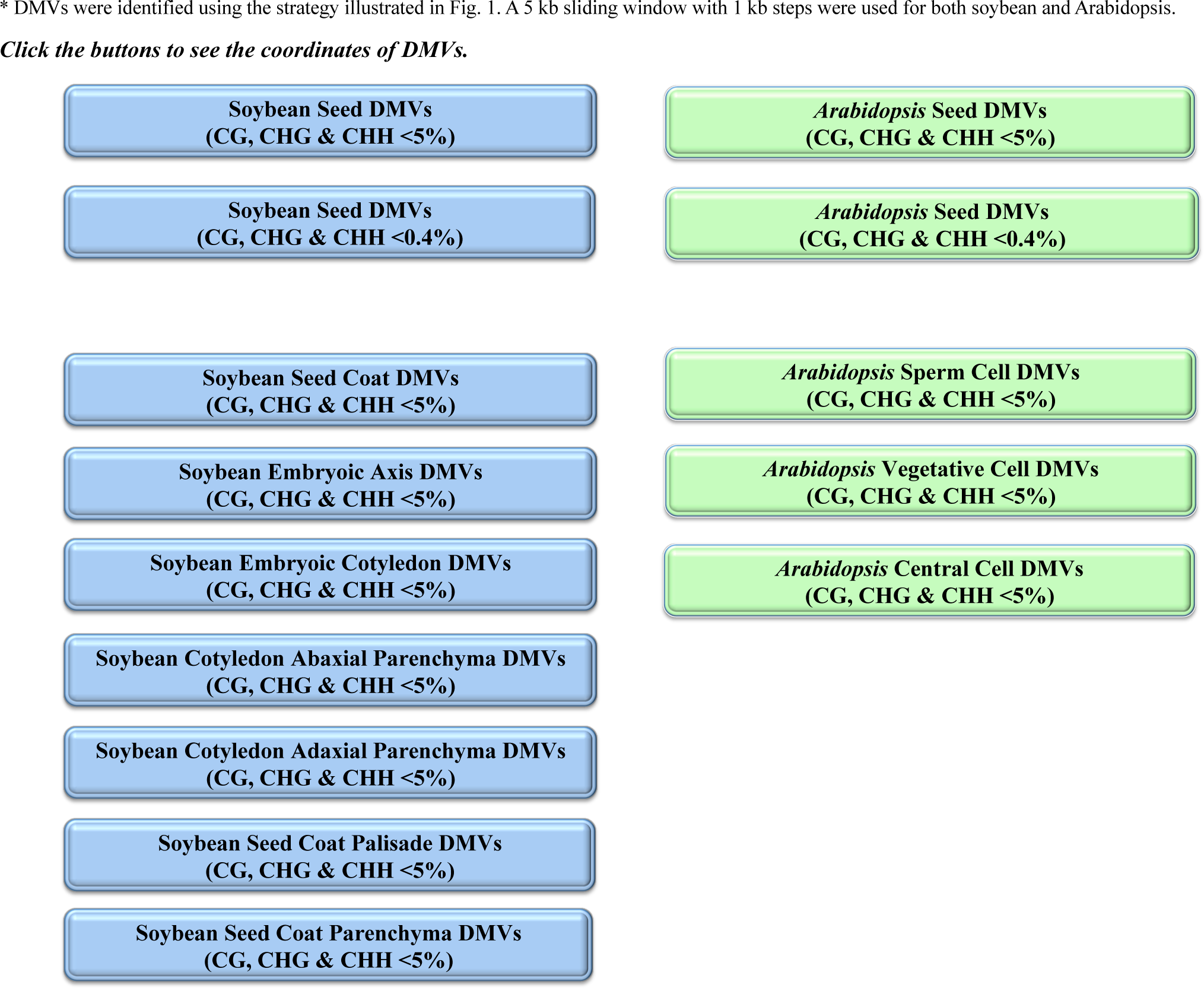
Soybean and *Arabidopsis* DMVs.

**Dataset S2.**
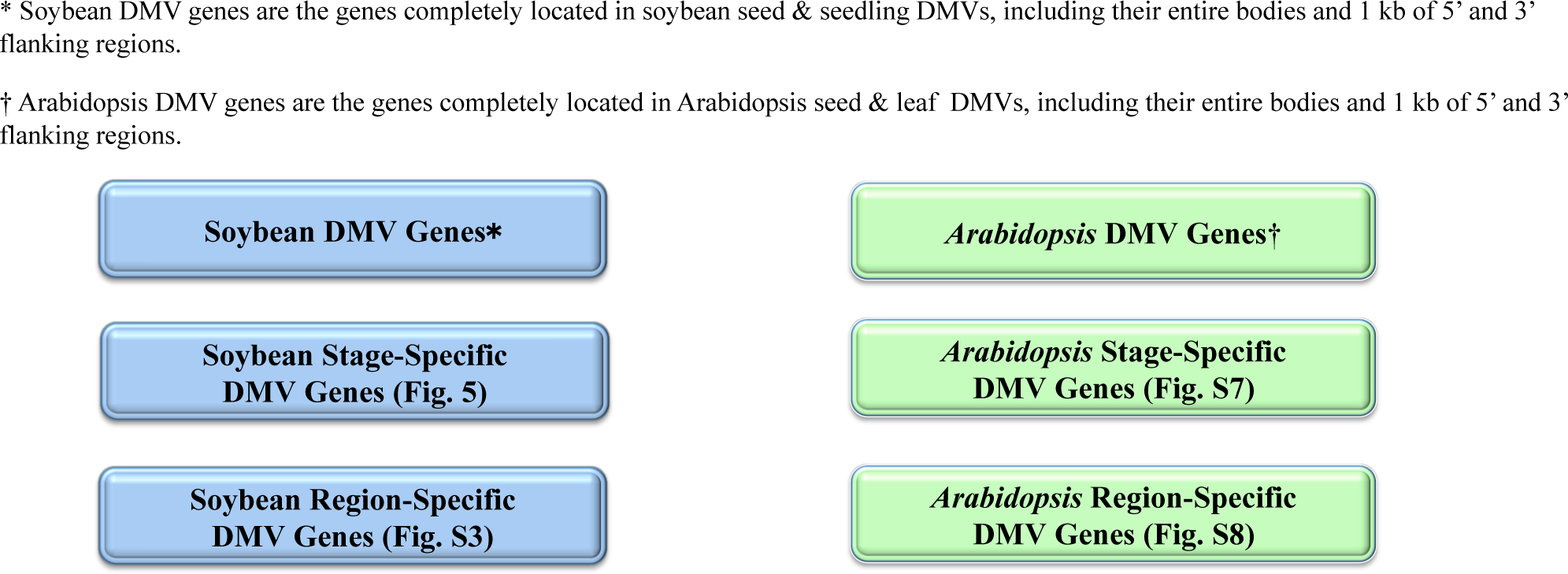
Soybean and *Arabidopsis* DMV Genes

**Dataset S3.**
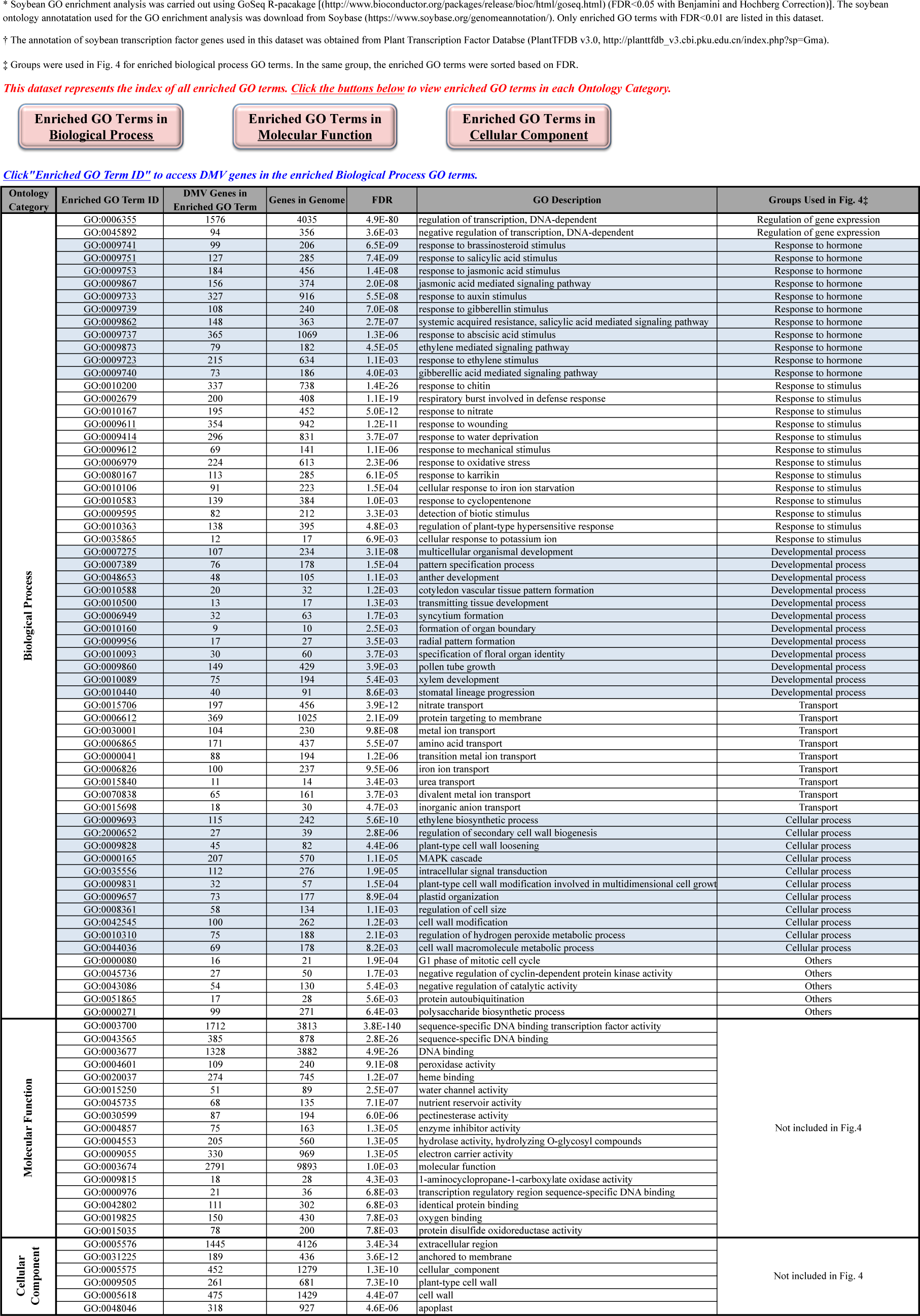
GO Enrichment Analysis* of Soybean DMV Genes, Including Transcription Factor Genes† (TFs)

**Dataset S4.**
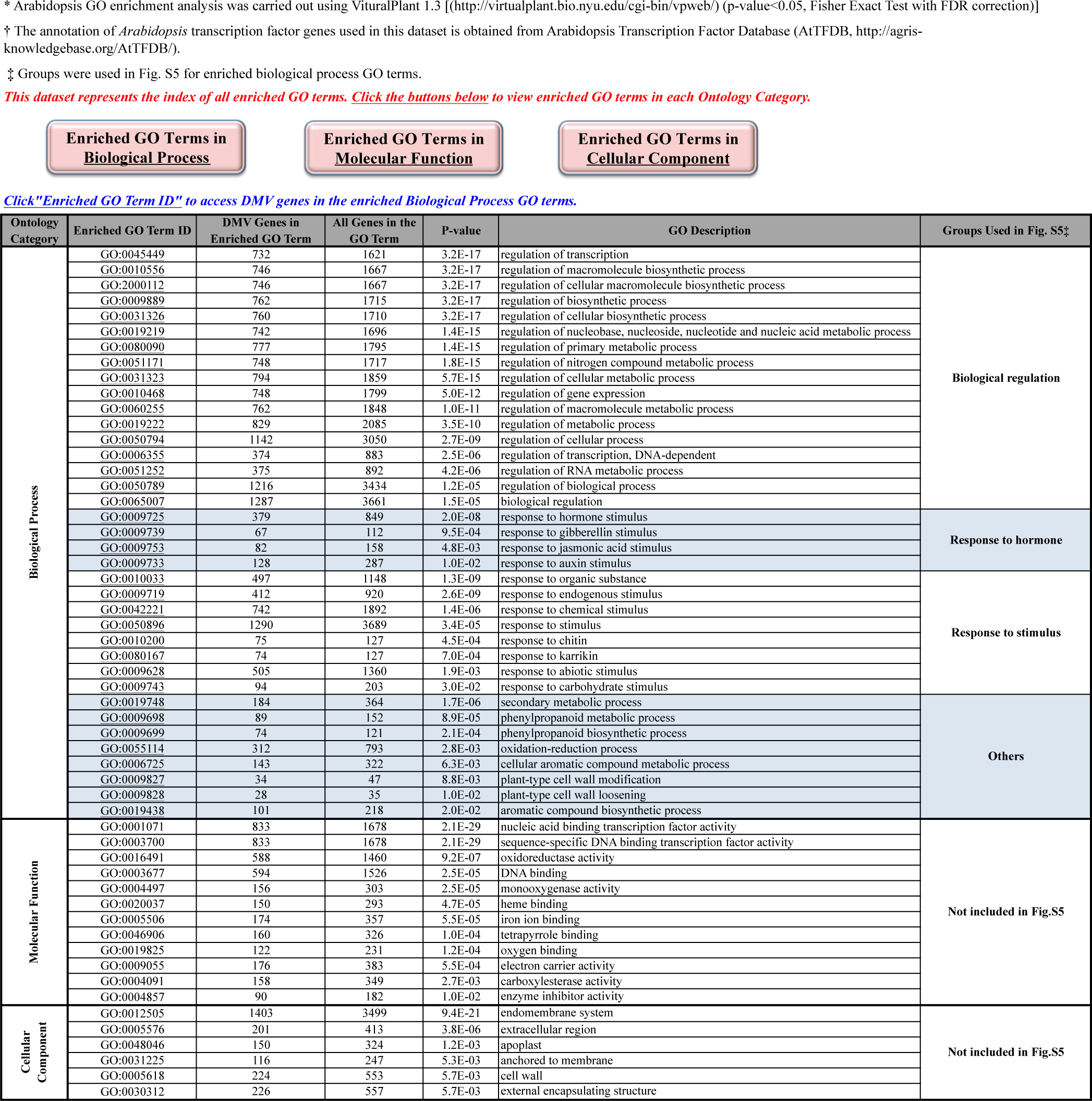
GO Enrichment Analysis* **of *Arabidopsis* DMV Genes, Including Transcription Factor Genes† (TFs)**

**Dataset S5.**
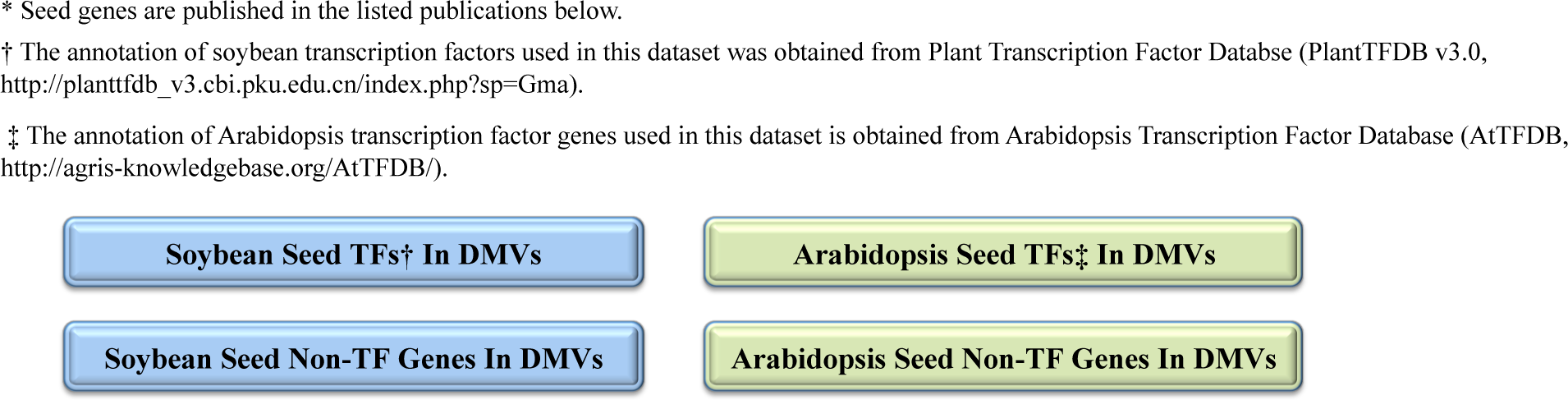
Soybean and *Arabidopsis* Seed Genes*

## Author contributions

M.C., J.-Y.L., M.P., J.J.H., and R.B.G. designed research; M.C., J.-Y.L., J.H., J.M.P., and R.B. performed research; M.C., J.-Y.L., J.M.P., and R.B. analyzed data; R.B.G. wrote the paper; J.J.H. and M.P. provided comments on the manuscript.

